# Precision Discovery of Novel Inhibitors of Cancer Target HsMetAP1 from Vast Metagenomic Diversity

**DOI:** 10.1101/2022.06.11.495772

**Authors:** Oliver W. Liu, Scott Akers, Gabriella Alvarez, Stephanie Brown, Wenlong Cai, Zachary Charlop-Powers, Kevin Crispell, Tom H. Eyles, Sangita Ganesh, Ee-Been Goh, Peter M. Haverty, William W. Hwang, Matthew Jamison, John L. Kulp, John L. Kulp, Zachary Kurtz, Andrea Lubbe, Aleksandr Milshteyn, Parisa Mokthari, Stephen G. Naylor, Samuel Oteng-Pabi, Ross Overacker, Andrew W. Robertson, Helen van Aggelen, Usha Viswanathan, Xiao Yang, Sam Yoder, Steven L. Colletti, Devin R. Scannell

## Abstract

Microbial natural products have long been a rich source of human therapeutics. While the chemical diversity encoded in the genomes of microbes is large, this modality has waned as fermentation-based discovery methods have suffered from rediscovery, inefficient scaling, and incompatibility with target-based discovery paradigms. Here, we leverage a metagenomic partitioning strategy to sequence soil microbiomes at unprecedented depth and quality. We then couple these data with target-focused, *in silico* search strategies and synthetic biology to discover multiple novel natural product inhibitors of human methionine aminopeptidase-1 (HsMetAP1), a validated oncology target. For one of these, metapeptin B, we demonstrate sub-micromolar potency, strong selectivity for HsMetAP1 over HsMetAP2 and elucidate structure-activity relationships. Our approach overcomes challenges of traditional natural product methods, accesses vast, untapped chemical diversity in uncultured microbes, and demonstrates computationally-enabled precision mining of modulators of human proteins.

## INTRODUCTION

Many of the most important therapeutic targets for cancer and other diseases are intracellular proteins ^1,2^. Despite the rise of mAbs and new therapeutic modalities ^3^, small molecule drugs remain best-suited for intracellular targets, offering cell permeability, oral administration and reliable delivery to diverse organs. However, many targets have proven very challenging for traditional synthetic chemistry which is biased to simpler SP2 chemistry, planar structures and further limited by Lipinski’s Rule-of-5 ^4^. In contrast, small molecule natural products (NPs), which are specialized metabolites encoded by biosynthetic gene clusters (BGCs) in the genomes of bacteria, fungi, and plants ^5^, are generally enriched for chiral centers, SP3 carbon atoms and rich 3D structures that enable them to mediate more complex protein interactions ^6,7^ NPs leverage enzymes that catalyze reactions that are often difficult to replicate via synthetic chemistry, have been selected to cross cell membranes, and typically violate Lipinski’s Rule-of-5 ^8,9^. Indeed, the remarkable properties encapsulated in these molecules is a direct consequence of billions of years of evolution to modulate cellular proteins and pathways ^10^.

While over half of approved small molecule drugs have NP origins ^5^, NPs have ceased to be a focus for large pharma. Traditional fermentation-based NP discovery approaches are plagued by high rates of rediscovery, inefficient scaling, and poor fit with modern discovery paradigms ^11,12^. A central limitation is that only ~1% of microbes are readily cultured in the lab and, therefore, amenable to fermentation-based discovery ^13 14^. Metagenomic approaches, which involve the direct capture of environmental DNA (eDNA) without culturing ^15^, have promised to eliminate the constraints imposed by culturing and fermentation-based discovery and to expand the potential for discovery by orders of magnitude ^16^. However, while congeners of known NP scaffolds have been discovered using metagenomics ^17–20^, de novo scaffolds with novel bioactivities have remained elusive outside of a few examples ^21^.

A major challenge for metagenomics has been that next-gen sequencing of complex metagenomic samples has failed to give the expected visibility into untapped BGC repertoires. The immense size of soil metagenomes, estimated to contain 10^4^-10^6^ unique phylotypes per gram, has proven too complex for shotgun sequencing. Short-read assemblies are too fragmented for the discovery of BGCs which range in size from 10 to 100+kb (Figure 1a) ^22^. Long-read technologies still lack the throughput needed for complex samples and/or suffer from high error rates ^23,24^. As a result, the vast majority of BGCs in these samples have remained undocumented.

**Figure 1.**
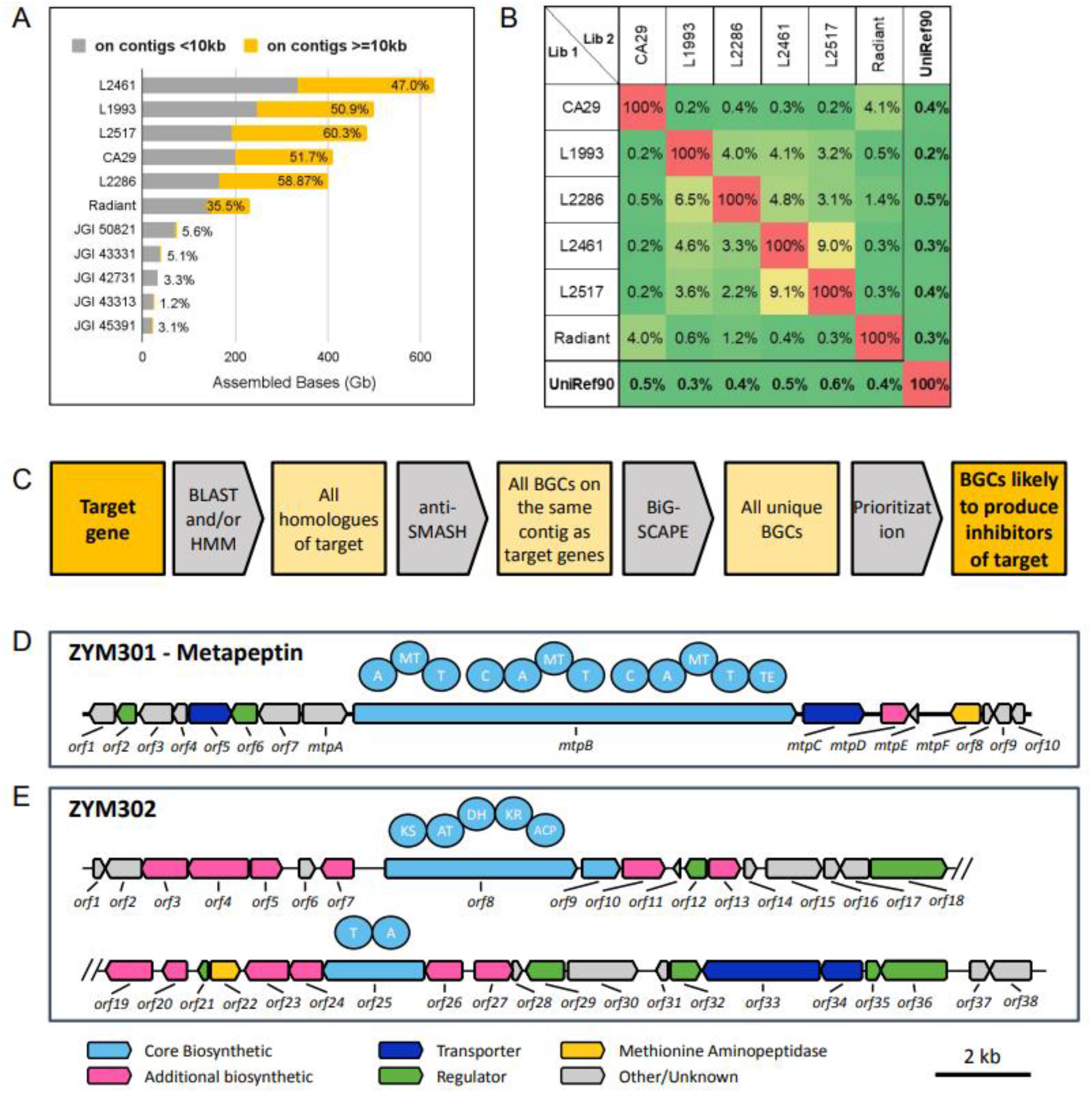
Metagenomic libraries contain vast, orthogonal diversity that can be searched using SREs. **(A)** Total assembly lengths for the 6 soil metagenomic libraries described in this paper and the 5 largest soil metagenomics datasets from JGI. Percentages indicate the fraction of assembled sequence contained on contigs >10kb. (**B)** Pairwise comparison of overlap between protein coding sequences in metagenomic libraries clustered at 90% aa identity as well as UniRef90 (in bold). (**C)** A flowchart showing the generalized workflow for identifying clusters that will produce inhibitors of target genes of interest. (**D-E)** ZYM301 (metapeptin) and ZYM302 gene clusters with color-coded gene function. Domain organization of PKS and NRPS-like genes are shown above the genes, labeled as adenylation (A), methyltransferase (MT), thiolation (T), condensation (C), thioesterase (TE), ketosynthase (KS), acyltransferase (AT), dehydratase (DH), ketoreductase (KR), or acyl carrier protein (ACP).

In order for a metagenomic NP discovery platform to truly yield novel, drug-like molecules, we reasoned that the platform needs 1) higher-quality metagenomic data that enables BGC discovery, 2) a computational approach to subsequently parse through millions of completely novel BGCs to identify those with therapeutically-relevant bioactivities, and 3) capabilities to produce and purify the bioactive molecules encoded by the selected BGCs.

In this paper, we present an end-to-end NP discovery platform that enables efficient discovery of novel NP modulators of proteins of interest from vast metagenomic diversity. Specifically, we discover novel inhibitors of human methionine aminopeptidase-1 (HsMetAP1), a key translational and cell cycle regulator implicated in multiple solid cancers ^1,25^ We detail efforts to sequence, assemble and catalog soil bacterial metagenomic diversity and biosynthetic potential on an unprecedented scale. From a database of millions of BGCs, we utilize self-resistance enzymes (SREs) to predict 35 to encode distinct, novel NP inhibitors of HsMetAP1, including two selected for experimental confirmation. Downstream technologies for heterologous expression, untargeted discovery of novel metabolites, and assignment of bioactivity to specific metabolites enabled the production, identification, and isolation of encoded molecules that validate our functional predictions. We further elucidated a novel, selective cyclic depsipeptide inhibitor of HsMetAP1, which we call metapeptin B.

## RESULTS

### Sequencing of large-insert cosmid metagenomic libraries enables access to massive metagenomic diversity including millions of novel BGCs

To assess the state of the art in soil metagenome sequencing, we analyzed the five largest soil metagenome datasets generated by the Joint Genome Institute (JGI) (https://img.jgi.doe.gov/cgi-bin/m/main.cgi) ^26^. These datasets, which were shotgun sequenced, range from 73.0Gb to 22.5Gb of assembled sequence; however, on average, only 3.7% (1.2%-5.6% range) of the assembled sequence in these datasets are found on contigs >10kb, indicating that the vast majority of these data are not useful for BGC discovery (Figure 1a).

We hypothesized that we could generate higher quality soil metagenomic assemblies by first partitioning the sample into smaller, lower-complexity sub-pools. Focusing on soil samples from 6 US locations, (Figure S1a), we built large-insert (35-40kb) cosmid libraries from eDNA, arrayed these into sub-pools of 6,000-25,000 cosmids and sequenced to 25X coverage (Illumina short-reads). The reduced complexity (from 10^4^-10^6^ to 50-250 bacterial genome equivalents per sub-pool), resulted in ~444Gb of assembled sequence per library with 229Gb (~51%) on contigs >10kb (Figure 1a). In comparison to the JGI datasets, we produced, on average, ~11.6X the amount of assembled sequence per library (444Gb vs 38Gb) with >140X more assembled data contained on contigs >10kb (229Gb vs <2Gb). Across the six libraries, we generated ~1.4Tb on assemblies >10kb, vastly expanding the potential for BGC discovery.

In order to assess the relative diversity found in each library, we annotated open reading frames in each library and compared the protein content (de-replicated at 90% amino acid identity) of the six libraries to each other as well as to the UniRef90 dataset. On average, each complete library contained 151M protein coding sequences, which is approximately the size of the UniRef90 dataset (release 2022_02). We saw <0.6% overlap between any of the libraries with UniRef90, consistent with the idea that the vast majority of soil microbial diversity has never been cultured and would not be found in public databases populated primarily with cultured organisms (Figure 1b). Strikingly, we saw only a 2.4% average overlap across all pairwise combinations of metagenomic libraries suggesting that the microbial diversity in each soil sample was largely orthogonal to the other samples. In total, across the six libraries, we annotated 903.2M open reading frames (de-replicated at 90% amino acid identity by library).

We used antiSMASH ^27^ to predict a total of >6.8M BGCs in the six sequenced libraries (Figure S1b). Of these, NRPS systems are most common (~2.5M), followed by terpenes (~1.3M), RiPP’s (~1.2M), and polyketides. While these are non-deduplicated BGC counts, based on the limited overlap in protein diversity between the six libraries, we anticipate that these data contain millions of distinct and novel BGCs. To our knowledge, this is the single largest database of BGCs globally. For comparison, a recent analysis of ~170,000 genomes in NCBI RefSeq database and ~47,000 metagenome assembled genomes from various sources identified a combined total of ~1.2M non-deduplicated BGCs ^28^.

### A resistance gene-based search strategy can be used to rapidly search metagenomic diversity for BGCs encoding predicted bioactivities of interest

To prioritize BGCs with potential therapeutic activity, we developed computational approaches to identify self-resistance enzymes (SREs) (Figure 1c) ^29^. An SRE, which typically is located within a BGC, encodes a resistant copy of the essential enzyme targeted by the NP produced by that BGC (e.g. resistant protease within BGC encoding a protease inhibitor). As a result, the function of an SRE effectively reveals the function of the BGC ^30,31^. While “resistant copy” SREs are rare ^29^, we predicted that with a database as large as ours, we could use this as a general method to connect novel BGCs to targets of interest.

We selected HsMetAP1 as a test. HsMetAP1 cleaves the N-terminal methionine residues of nascent peptides and is important for cell cycle regulation. It re-emerged as a priority for several solid tumors in a recent large-scale CRISPRi screen ^1,25^ There is one known class of bacterial NP inhibitors of HsMetAP1, called the bengamides, that in *Myxococcus viriescens*, are encoded by a BGC that contains a methionine peptidase SRE ^32^.

We searched a subset of our metagenomic database (~1.2M BGCs) for BGCs containing a MetAP1 homolog within the cluster. The resulting BGCs were de-replicated and computationally prioritized (Figure 1c) to identify putative SREs. In total, we identified 35 unrelated BGCs that met our criteria (Table S1). Notably, none of the identified BGCs resemble any characterized biosynthetic systems found in the MiBIG database ^33^ including that for bengamide, highlighting the novelty of metagenomic diversity.

### Heterologous expression of BGCs containing putative MetAP1 resistance genes produces lysates with predicted inhibitory bioactivity

We selected two of these BGCs for expression studies to validate our resistance gene-based functional predictions. ZYM301, contains genes encoding for an NRPS (*mtpB*), a methyltransferase (*mtpD*), a transporter gene (*mtpC*), two genes of unknown function (*mtpA* and *mtpE*), and the methionine aminopeptidase (*mtpF*) (Figure 1d; Table S2). Based on domain structure, *mtpB* is predicted to incorporate N-methyl-L-tyrosine, N-methyl-L-threonine, and N-methyl-L-valine residues. ZYM302, contains a single-module PKS (*orf8*), an NRPS-like gene (*orf25*), thirteen biosynthetic genes, two transporters, seven regulators and the methionine aminopeptidase (*orf22*) (Figure 1e; Table S3). Little can be predicted about the biosynthesis of ZYM302 metabolites.

ZYM301 and ZYM302 BGCs were conjugated into Streptomyces albus J1074 and the resulting exconjugates were fermented in mO42 media and crude organic extracts. Using an established colorimetric assay we detected HsMetAP1 inhibitory activity in both extracts relative to empty vector controls (Figure 2a), validating our SRE-based BGC selection strategy.

**Figure 2.**
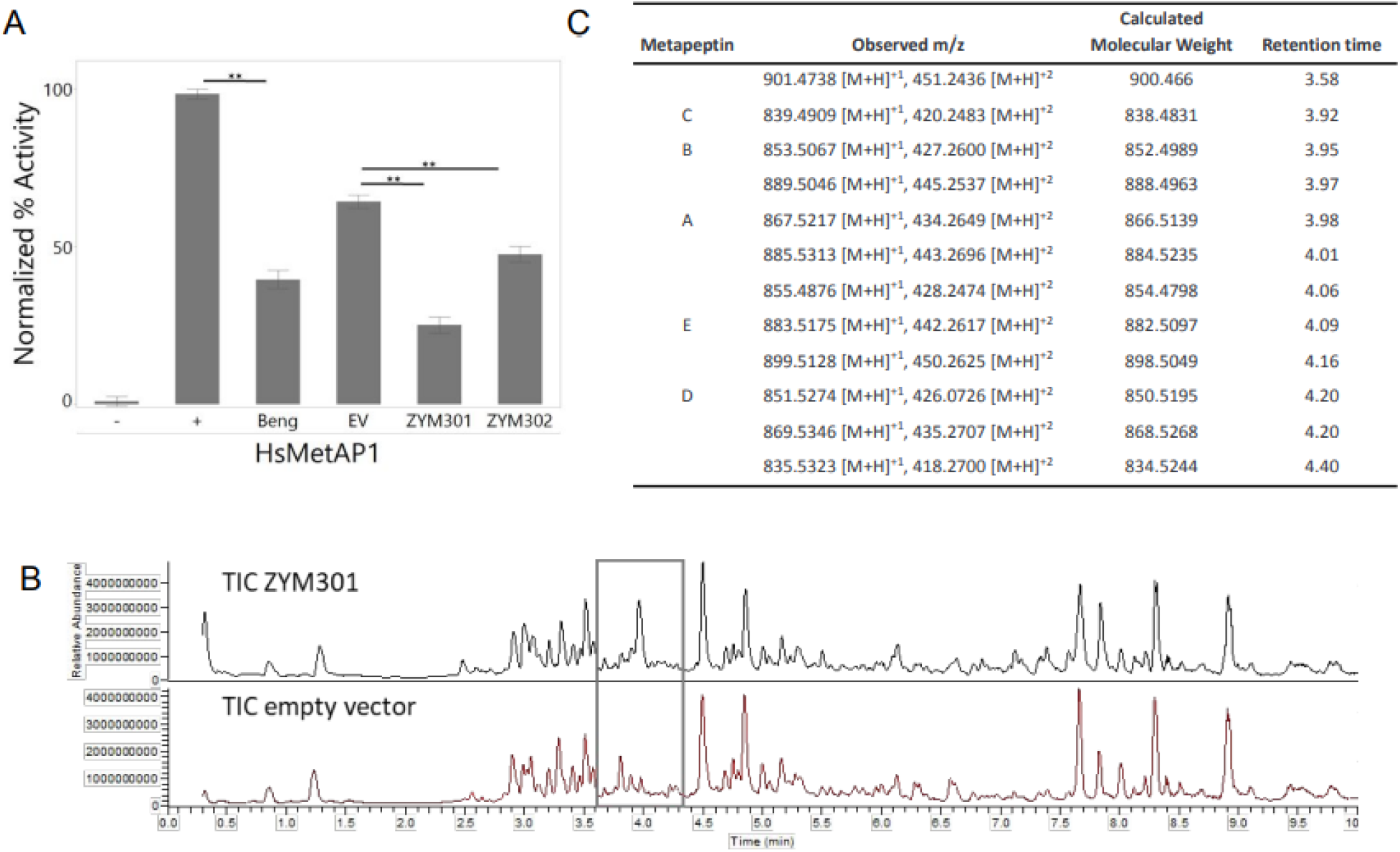
Crude extracts from *S.albus:ZYM301* and *S.albus:ZYM302* inhibit HsMetAP1. **(A)** Crude extracts (0.5mg/ml) from S. albus:ZYM301, *S. albus:ZYM302, S. albus* empty vector control (EV) and bengamide B (Beng) control (100 μM) were assayed for HsMetAP1 inhibitory activity. Controls included a DMSO vehicle control (+enzyme) and reactions that did not contain hsMetAP1 (−enzyme). Error bars represent the standard deviation of the mean, n=3. **Statistical significance assessed via Dunnett’s, p-value (<0.001). **(B)** Total Ion Chromatograms of a representative *S. albus:ZYM301* and empty vector control sample. Box highlights the region where novel features were detected. (C) Novel compounds produced by *S. albus:ZYM301* that were identified by untargeted LC-MS/MS analysis.

LC-MS/MS analysis of *S.albus:ZYM301* identified one major novel species (m/z 867.5217), which we call metapeptin A, and 11 minor species that include metapeptins B-E (Figure 2b-c, Figure S2). All had a common MS2 fragment of m/z 164.1082, suggesting a N-methylated tyrosine residue (Figure S3). Molecular weight ranges from 834 to 898 Da indicate that each amino acid was incorporated multiple times. Three compounds had differences of m/z 14.015, suggesting differential methylation. For *S.albus:*ZYM302, differential analysis yielded 9 features but no MS2 similarities (data not shown). All compounds/features detected for ZYM301 and ZYM302 were determined to be novel but ZYM301 was prioritized for further exploration due to higher activity in initial assays (Figure 2a), higher titers and robust MS profile.

To determine which metabolite(s) produced by *S. albus:ZYM301* is responsible for HsMetAP1 inhibitory activity, we opted for a biochemometric approach to statistically link novel compounds to bioactivity ^34,35^. Three parallel fractionation strategies (normal phase, reverse phase, and size exclusion) were pursued, each generating seven fractions (total of 21) which were subjected to bioassay and metabolomics analyses (Figure 3a). Partial least squares regression was used to calculate selectivity ratios ^36^ that were projected onto a molecular network of all MS2 features (Figure 3b). The six features with the highest ratios were all associated specifically with metapeptin B (Figure 3c), accounting for most if not all observed bioactivity.

**Figure 3.**
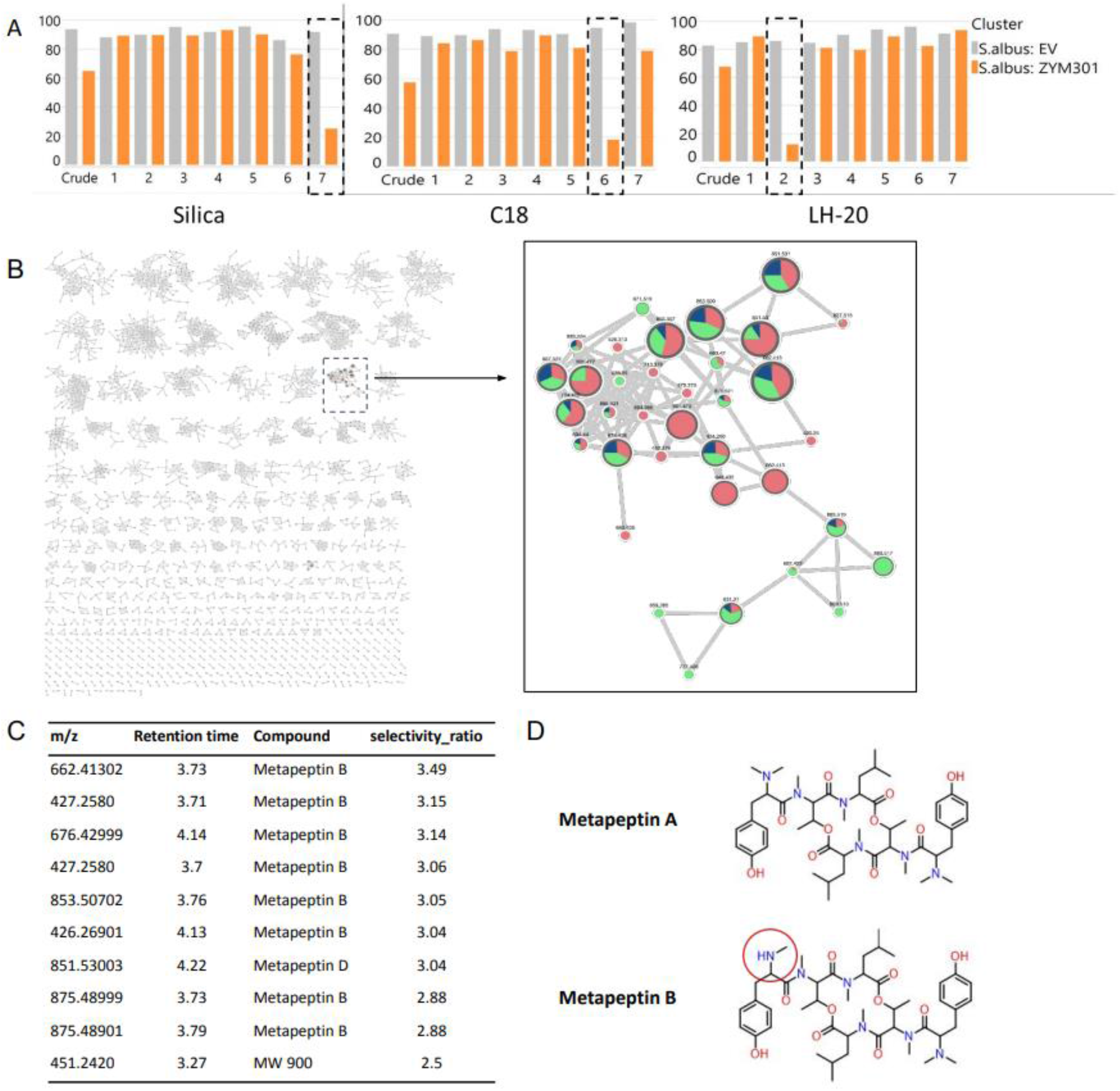
Biochemometric strategy indicate metapeptin B is largely responsible for observed bioactivity. **(A)** 21 fractions generated using C18, Silica and LH20 size exclusion chromatography columns were assayed for HsMetAP1 inhibitory activity. Dotted boxes outline significant inhibition. **(B)** A molecular network of all the features detected with MS2 data across 21 fractions, and a magnified view on the connected component that contains the compounds with the highest selectivity ratio. The node size indicates the selectivity ratio and the pie chart indicates the fractionation method it was observed in (green: Silica, red: C18, blue: LH20). **(C)** Top 10 differentially expressed features sorted by selectivity ratio and the compounds with which they are associated. **(D)** Structures of metapeptin A and B. The red circle highlights the single methyl difference between the two molecules.

### Metapeptin B is a novel cyclic depsipeptide that differs from metapeptin A by only a single methylation

Given higher production titers, we first elucidated the structure of metapeptin A (Figure 3d). The ion peak at m/z of 867.5217 suggested a molecular formula of C_46_H_70_N_6_O_10_ while analysis of 2D NMR spectra, demonstrated the amino acid components as *N,N*-dimethyl tyrosine, *N*-methyl threonine and *N*-methyl leucine, consistent with our bioinformatic prediction. Heteronuclear multiple bond correlation (HMBC) spectra established the backbone of the 14-member cyclic peptide (Table S5, Figure S4, Supplementary Text).

The formula C_45_H_68_N_6_O_10_ (m/z 853.5067) indicated that metapeptin B and A differ by a single CH_2_. Under optimized conditions, metapeptin B produced an extra fragment pair with m/z 164.1070 and 150.0913 linking the missing CH2 to the *N, N*-dimethyl-Tyr moiety (Figure S5). While further MS data could not definitively confirm which carbon is missing, we propose that metapeptin B is an *N*-mono-methylated congener of metapeptin A (Figure 3c). First, *mtpB* is bioinformatically predicted to utilize tyrosine, making the incorporation of *N, N*-dimethyl-hydroxyphenylglycine highly unlikely. Second, another leaky NRPS-embedded *N*-methyltransferase domain is known which produces demethylated shunt products at reduced titers similar to the ~22 fold lower yield of metapeptin B vs. A ^37^. The structures of metapeptin A and B have never previously been observed, consistent with the novelty expected from metagenomically-sourced BGCs.

### Metapeptin B is a sub-micromolar inhibitor of EcMetAP and is highly selective for HsMetAP1 over HsMetAP2

We conducted a series of enzyme assays with purified metapeptin A and B. As predicted by our biochemometric analysis, metapeptin B inhibited HsMetAP1 with an IC_50_ of ~50uM while metapeptin A showed no activity even at the highest concentration tested (2mM) (Figure 4a).

**Figure 4.**
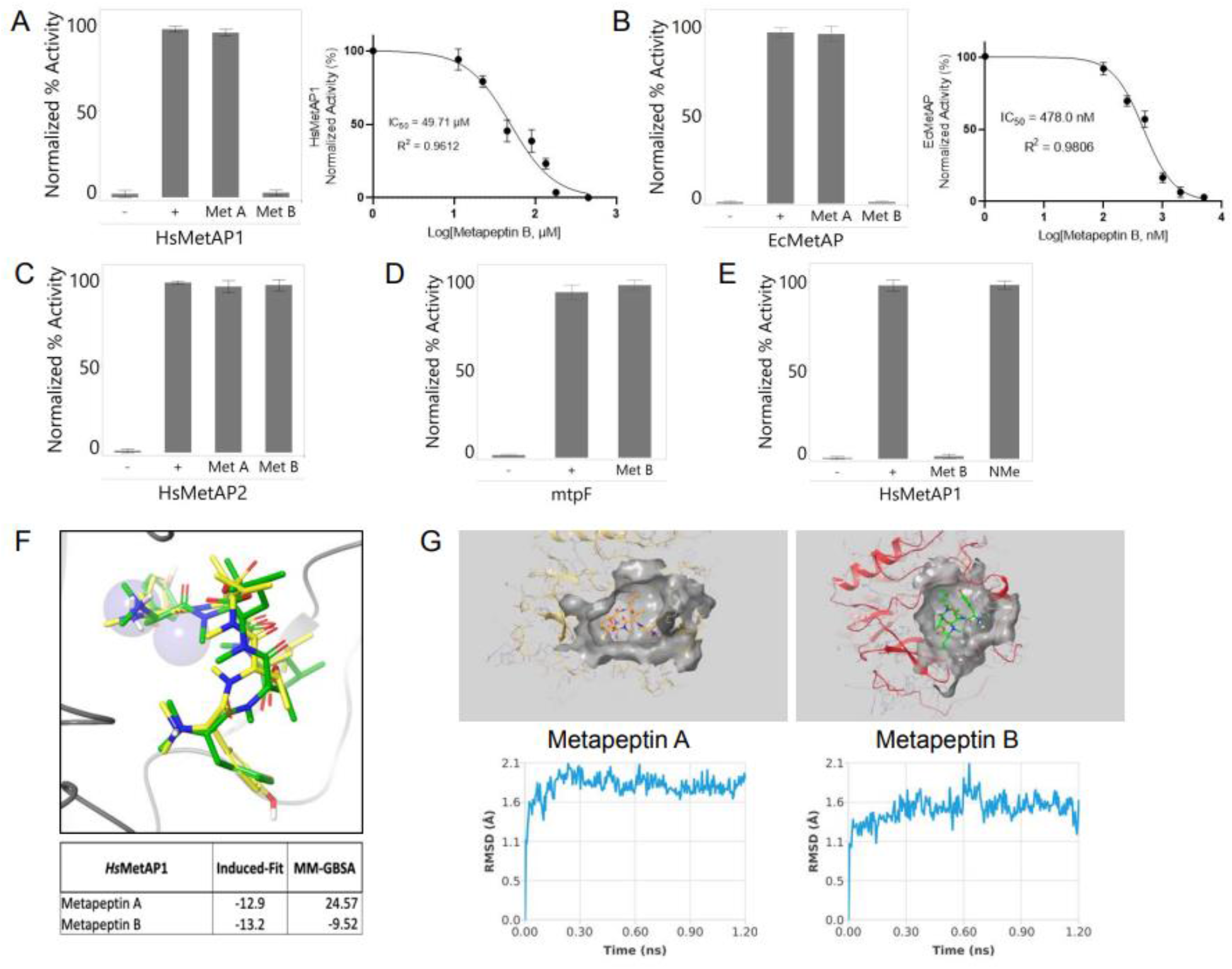
Metapeptin B encodes a selective inhibitor of HsMetAP1. Metapeptin A (MetA) and metapeptin B (MetB) (100 μM) were assayed for inhibitory activity against **(A)** HsMetAP1, **(B)** E. coli MetAP (EcMetAP), and **(C)** HsMetAP2. Error bars represent the standard deviation of the mean, n=3. **(A-B)** Dose response curve of Metapeptin B (0.5 μM – 500 μM) against hsMetAP1 (0.5 μM – 500 μM) and Ec.MetAP1 (100nM - 5,000nM). Non-linear regression (variable slope) analysis used to fit the curve. Error bars represent the standard deviation of the mean, n=9. Statistical significance assessed via ANOVA, p-value (<0.001). For all enzymatic assay, controls include a DMSO vehicle control (+enzyme) and reactions that did not contain enzyme (−enzyme). (D) Metapeptin B was assayed for mtpF (putative resistance enzyme) inhibitory activity. Error bars represent the standard deviation of the mean, n=3. (E) Metapeptin B and the NMe-Monomer linear tripeptide (100μM) were assayed for HsMetAP1 inhibitory activity. (F) Induced fit docking shows conformational differences resulting in better docking score and MMGBSA energies (more negative is more favorable) for metapeptin B including a large DDGs solvation change. (G) Lower RMSD for distance to binding pocket residues for metapeptin B indicates more stable binding.

Similarly, metapeptin B, but not metapeptin A, inhibited the *E. coli* methionine aminopeptidase homolog (EcMetAP) (Figure 4b). Notably, metapeptin B activity against EcMetAP was ~100-fold stronger (IC_50_=~500nM) than against HsMetAP1 (Figure 4a-b). Both results highlight the importance of the single methyl difference between metapeptins A and B. We saw no inhibition of the other methionine peptidase in the human genome, HsMetAP2, by metapeptin B at 2mM (Figure 4c), indicating metapeptin B is highly selective for HsMetAP1 over HsMetAP2.

To confirm that the putative SRE gene in the metapeptin BGC (*mtpF*) functions as a resistance gene, we tested the ability of metapeptin B to inhibit the activity of the methionine peptidase variant encoded by *mtpF*. Consistent with its predicted role in alleviating metapeptin B toxicity, we found that metapeptin B had no effect on the activity of the methionine peptidase encoded by *mtpF* at the highest concentrations tested (200mM) (Figure 4d)

### Asymmetric methylation of metapeptin B stabilizes the interaction of the cyclized dimer with HsMetAP1

To understand how a single methyl difference between metapeptins A and B drives such major differences in activity, we conducted a series of cheminformatic analyses (Figure 4f-g). Induced-fit docking on HsMetAP1 (PDB:6LZC) showed that differences in the methyl amine sites and a flipping of the methyl amide linkers (Figure 4f) contribute to metapeptin B having a significantly lower MMGBSA energy compared to metapeptin A. In addition, metapeptin B has a stronger electrostatic interaction with the positively charged metal ions due to a more favorable orientation (Figure S7), stronger dipole (metapeptin A: 7.05D versus metapeptin B: 7.34D) and less positively charged surface area (pKa of metapeptinA (7.76) versus metapeptinB (7.46)). Finally, molecular dynamic simulations showed a lower RMSD to the H212 residue for monomethyl amine of metapeptin B compared to the dimethyl amine of metapeptin A (Figure 4g).

Lastly, to understand the importance of the asymmetric ring structure to these differences, we designed and synthesized the singly methylated tripeptide monomer (NMe-Monomer) (Figure S6). The linear NMe-Monomer demonstrated no detectable inhibition at 2mM, confirming the cyclization of metapeptin B is critical for its target engagement, most likely due to conformational rigidity (Figure 4e).

## DISCUSSION

In this paper, we identified millions of previously unexplored BGCs by creating high quality assemblies from diverse soil metagenomes. We demonstrated an efficient computational search that allowed us to down-select from >1M BGCs to 35 candidates predicted to encode novel methionine peptidase inhibitors. These BGCs encode diverse chemistries (Table S1) and are unrelated to characterized BGCs in the MiBIG database. We validated the predicted activity for ZYM301 (NRPS) and ZYM302 (NRPS-like) and ZYM301 was found to encode metapeptin B, a novel depsipeptide inhibitor. Most importantly, our approach surmounts the key challenges of traditional fermentation-based NP discovery (e.g., rediscovery, phenotypic rather than target-based screening and poor scalability) ^11,12^ and thus opens up the massive chemical diversity in nature for modern drug discovery.

Scale is a key consideration for our approach. In the case of methionine peptidases, we estimate that no more than 1 in 20,000-80,000 BGCs would meet our criteria for further exploration. While this number will no doubt vary by target family, it implies millions of BGCs are required to confidently pursue a drug discovery program where the goal is to identify multiple diverse, high-quality leads for optimization. By contrast, only one class of bacterial NP HsMetAP1 inhibitors, the bengamides, was previously known ^38^. Our ability to identify up to 35 additional diverse BGCs encoding HsMetAP1 inhibitors suggests that much more robust NP drug discovery programs are possible with sufficiently large databases.

Our data also indicate that unexplored metagenomic diversity is extremely large. Despite finding potentially many BGCs that encode HsMetAP1 inhibitors, we found both ZYM301 and ZYM302 only once each. We failed to find any members of the bengamide family. Together with the very low overlap between our six metagenomic samples (average ~2.4% of proteins shared between pairs of libraries), these data suggest that even with the massive expansion of access described here, we are just beginning to explore metagenomic diversity. This contrasts sharply with the high rediscovery rates of fermentation-based NP discovery and highlights the profound limitations of strain collections and focusing on the cultured space for innovative drug discovery.

While the strategy described here relies on the identification of bacterial NPs that inhibit bacterial enzymes, we believe our approach has broad applicability to human targets. First, there is abundant precedent for drugs derived from bacterial NPs being approved as modulators of human therapeutic targets (e.g., Kyprolis, based on epoxomicin and Rydapt, based on staurosporine). Second, the resistance-gene approach utilized in this study appears to be relatively insensitive to evolutionary distance. Methionine peptidases are only ~40% identical between humans and bacteria. Our approach also works for proteasome subunits (data not shown), the target of Kyprolis, which are only ~30% identical. Conserved pharmacological tractability despite low protein sequence identity presumably reflects the deep importance of core cellular biology. Lastly, we have built a similar fungal metagenomic database that we believe will expand the set of pharmacologically tractable human targets to include many without homologs in bacteria

The key constraint to our approach is the need to heterologously express BGCs to validate predicted activity and elucidate structures. In this paper, we conjugated ZYM301 and ZYM302 into *S. albus* without promoter refactoring. Both expressed sufficiently to confirm activity, but we focused on ZYM301 in part due to the low titers for ZYM302. While there are many levers to increase production (e.g., media and fermentation optimization), in our experience synthetic biology is key. Initial engineering can take the form of refactoring promoters within the BGC but at later stages, large-scale, genome-wide host engineering may also be required. Indeed, expressing BGCs at a scale (both throughput and yield) that is matched to the needs of drug discovery is, in our view, the key to success with this approach.

In summary, our approach to NP discovery allows us to efficiently leverage billions of years of evolution for modern drug discovery. By combining metagenomics, data science and synthetic biology we demonstrate a scalable platform for precision discovery of NPs with activity against targets of interest. Moreover, our data support the view that metagenomic diversity is extremely large ^39^, and that the universe of NPs relevant for targeted human drug discovery is larger than previously imagined.

## MATERIALS AND METHODS

### eDNA libraries construction and sequencing

Soil eDNA libraries were constructed following a protocol adapted from the Brady protocol ^40^. Briefly, eDNA was released from ~125g to 500g of soil sifted through a 2mm sieve by suspending it in an SDS detergent buffer and heating the mixture at 70°C for 2 hours. After removing debris by centrifugation, DNA was extracted by alcohol or PEG precipitation and further purified and size selected by field inversion gel electrophoresis. DNA in the 35-40kb fraction was end-repaired and ligated into an appropriate library vector. Cosmid libraries were constructed using MaxPlax™ Lambda Packaging Extracts (Lucigen Corp, Middleton, WI).

Prior to preparing sequencing libraries the cosmid backbone was removed by digesting each DNA sub-pool with an appropriate restriction enzyme and removing the small backbone fragments. Purified insert sub-pools were then tagmented and barcoded using unique combinations of Illumina i5 and i7 index primers. Final barcoded DNA libraries were sequenced on an Illumina Novaseq instrument.

### Sequence Annotation

Following assembly, using metaSPAdes ^41^, open reading frames (ORFs) were called using Prodigal ^42^and taxonomic classification for each contig was assigned using Kaiju ^43^. ORFs were further annotated with PFAM domains using hmmer ^44^ and PFAM-A database ^45^. Biosynthetic gene clusters were predicted on all contigs using antiSMASH v5 ^27^.

### JGI analysis

To evaluate how Zymergen soil metagenomic libraries compare to the largest publicly available soil metagenomic datasets, we looked to the Joint Genome Institute’s Integrated Microbial Genomes & Microbiomes (IMG/M) database (https://img.jgi.doe.gov/cgi-bin/m/main.cgi). We identified the 5 largest datasets in IMG/M based on the assembled genome size, corresponding to IMG Genome IDs 3300050821, 3300043331, 3300042731, 3300043313, and 3300045391. The assembly lengths of all contigs larger or equal to 10kb were extracted from the coverage statistics file associated with each dataset and summed to yield the total assembly lengths contained on contigs >10kb.

### Library comparisons

To evaluate the overall novelty of sequence in the libraries, we first clustered each library independently at 90% amino acid sequence using MMseqs2 algorithm ^46^, such that the clusters were composed of sequences containing at least 11 residues and at least 80% overlap with the seed sequence of the cluster (*i.e*., the longest sequence in the cluster). Subsequently, the representative cluster outputs from individual libraries were co-clustered together with the UniRef90 dataset (Release: 2020_01).

### MetAP1 resistance gene search

To identify homologues of HsMetAP1, a BLASTp search was run against the Radiant and CA29 metagenomic libraries using the HsMetAP1 gene (P53582) as the query. To allow the search to include distant homologues, a permissive cut off of e-value < 10 was chosen. The resulting hits were narrowed down to those that share a contig with a BGC, as called by antiSMASH (Blin et al. 2021). The BGCs closely associated with the MetAP1 homologues were run through BigScape to group together similar clusters, allowing dereplication at a cutoff of 0.7 ^47^. Finally, the remaining BGCs were prioritized based on a number of factors including the similarity of the putative resistance gene to HsMetAP1, the distance of the putative resistance gene from the BGC, the presence of the resistance gene in an operon with a biosynthetic gene, and the predicted completeness of the cluster. This resulted in a list of 35 “high quality” clusters that were candidates for producing a inhibitor for methionine aminopeptidases.

### Isolation of cosmids from eDNA libraries

To isolate cosmids containing BGCs of interest, primers specific to the clusters of interest were designed to generate a 400-500kp amplicon. These primers were used to track the cosmids through multiple rounds of serial dilutions by PCR. Briefly, a library well containing the cosmid of interest was used to inoculate an overnight culture in LB supplemented with 100 μg/ml carbenicillin. Using OD600nm, approximately 30 cells/well were inoculated in a 384-deep well plate and grown overnight. Positive wells as assayed by PCR were then plated to single colonies on LB (100 μg/ml carbenicillin) Q-trays to select for colony-PCR positive clones. Isolated clones were sequence verified by Illumina NGS sequencing via tagmentation.

### Assembly of BGCs into heterologous expression vectors

For heterologous expression of ZYM301 and ZYM302 after isolating the cosmids containing the BGCs, yeast homologous recombination was used to transfer the BGCs to integrative (pTARw) expression vectors containing yeast replication origins and selection markers. Briefly, pTARw were digested with I-SceI and PacI to linearize and expose ends that have homology to cosmids sequences flanking the eDNA encoding ZYM301 and ZYM302 respectively. These digested vectors were co-transformed with the cosmids of interest into *Saccharomyces cerevisiae*(BMA64) using a standard LiAc/SS carrier DNA/PEG method ^48^. DNA from PCR-positive yeast colonies were isolated using the ChargeSwitch™ Plasmid Yeast Mini Kit (Invitrogen) and electroporated into epi300 electrocompetent cells. DNA extracted from positive colonies checked by cPCR was sequence verified by Illumina NGS sequencing via tagmentation.

### Heterologous expression

Heterologous expression plasmids for ZYM301 and ZYM302 were transformed into *Escherichia coli* S17.1 cells and transferred into *Streptomyces albus* via conjugation. Exconjugant colonies that grow on Mannitol Soya (MS) agar plates with nalidixic acid (30 μg/ml) and apramycin (50 μg/ml) were restreaked to single colonies. Four colonies that passed colony PCR verification were glycerol stocked and then used to inoculate triplicate 3mL seed cultures (starting OD450=0.05) in Tryptic Soy Broth (TSB) + apramycin (50 μg/ml) along with appropriate vector and media only controls. Seed cultures were grown for 3 days at 30°C and then diluted 1:10 in 4 different media (O42, mO42, R5A and ISP4) for fermentation. After 7 days of incubation at 30°C for 7 days the cultures were then extracted as described below for AC analysis.

### MetAP1 enzymatic assay validation

The primary methionine aminopeptidase colorimetric activity assay is based on a commercial kit (R&D Systems) and further developed and optimized as follows. The reaction occurs in two steps: enzyme activation and product detection. *Enzyme activation*, each well contains a 25 uL mixture of activation buffer: 50 mM HEPES, 0.1 mM CoCl2, 0.1 M NaCl, pH 7.5; 100 uM of fluorogenic tripeptide substrate Methionine-Glycine-Proline-7-amido-methylcoumarin (R&D Systems ES017); and 2 ug/mL of MetAP enzyme (R&D Systems 3537-ZN). *Product detection*,each reaction well is supplemented with 25 uL of 2 ng/mL DPPIV/CD26 (serine exopeptidase diluted in activation buffer).

The assay was performed at room temperature in an opaque 384-well polystyrene plate and measured in a black, flat bottom, 384-well plate. The assay is initiated as MetAP is mixed with the tripeptide substrate and incubated for 5 minutes.The initial reaction will result in the cleavage of methionine, yielding a dipeptide product, Gly-Pro-AMC. MetAP is then inactivated by heating the reaction to 100 °C for 5 min and cooled on ice for an additional 5 min. Detection of the dipeptide product is carried out via two orthogonal methods; degradation of the dipeptide product via DPPIV protease and targeted LC-MS analysis of the dipeptide product. Incubation of the reaction mixture with a DPPIV solution at room temperature for 10 minutes results in the hydrolysis of the dipeptide and release amido-methylcoumarin, which is measured at 380/460 nm excitation/emission on a Tecan Spark microplate reader. The release of AMC, measured via fluorescence, corresponds to the activity of the MetAP under analysis. Alternatively, the dipeptide-AMC product is measured via LC-MS, using a standard (Sigma Aldrich G2761) to quantify MetAP activity. Both analysis methods have been proven to yield comparable signals, validating either method as a tool characterizing enzyme activity.

### MetAP1 enzymatic assay background controls

As is typical in enzyme kinetics, the initial rate of enzyme hydrolysis is used to measure enzyme activity, with background hydrolysis being subtracting from the enzymatic output to observe accurate MetAP activity. Background controls were vetted by removing key reagents from the reaction mixture to ensure that activity was dependent on the expected agents (substrate, enzyme, co-factors) and when any one was not present, enzyme activity above background was not observed. All MetAP enzymes tested tolerated up to 10% DMSO and 5 % Methanol without significant enzyme inactivation.

Two additional controls were established to confirm that observed inhibition is specific to MetAP and not the DPPIV protease used in the colorimetric readout. First, bioactive fractions/molecules (including metapeptin B) were incubated with the dipeptide product and the DPPIV protease. In all cases, the fractions/molecules showed no inhibition of the hydrolysis of the dipeptide by the DPPIV protease and release of AMC, indicating that the observed inhibition is specific to MetAP. Second, to address concerns about potential false positives with the colorimetric assay, LC-MS analysis was also used to confirm the increase in the MetAP cleavage product, GP-AMC, in positive control reactions.

Finally, *S. albus* fermentation extracts can have overlap in fluorescence with the AMC readout used to measure activity in the colorimetric assay. To control for this potential interference, an inhibitor control was established by incubating the test inhibitor in the activation buffer and measuring its fluorescence. This value was then subtracted from the MetAP+inhibitor reaction to determine the true impact of the inhibitor on enzyme activity. Conversely, *S. albus* extracts may also cause non-specific inhibition at relatively high concentration within the assay. We determined that 0.1-0.5 mg/ml was an acceptable range for the working concentration of extracts/fractions that enable the detection of inhibition while maintaining relatively low background fluorescence.

### Sample Preparation for UPLC-MS/MS analysis

Three mL of fermentation broth was extracted twice with 3 mL ethyl acetate (HPLC grade, Fischer) by shaking for 1 minute at 1000 rpm followed by sonication for 15 minutes. Samples were centrifuged, and the organic layer was collected and pooled to yield 6 mL of extract. The ethyl acetate was removed under reduced pressure in a Speedvac Savant (Thermo). Dry samples were suspended in 120 uL methanol (LC-MS grade, Fisher) and transferred to HPLC vials. A pooled sample for each medium was generated by combining 30 uL aliquots from replicate samples of each medium type.

### UPLC-MS(/MS) data acquisition

Samples were subjected to ultra performance liquid chromatography mass spectrometry on a Q-Exactive Mass Spectrometer (Thermo Fisher) connected to a Vanquish Liquid Chromatography system (Thermo Fisher). A gradient of water (mobile phase A) and acetonitrile (mobile phase B), each containing 0.1% formic acid, was employed with a flow rate of 0.5 mL/min on a Zorbax Eclipse Plus C18 RRHD 2.1 × 50 mm, 1.8 μm column (Agilent), operated at 40C. The gradient started at 2% mobile phase B, holding for 1 min, followed by a linear gradient to 100% mobile phase B over 7 minutes, and then held at 100% mobile phase B for 2 minutes, returning to initial conditions over 0.1 min and holding for 0.9 min for a total run time of 11 min. The mass spectrometer was operated at spray voltage: 3.6 kV, capillary temperature: 275, sheath gas flow rate: 25, auxiliary gas flow rate: 10, S-lens RF level: 70. Full scan mass spectra were acquired in positive and negative ionization mode from m/z 200-1500 at 70K resolution and ACG target of 3e^6^, and maximum ion fill time of 200 ms. Data-dependent MS2 spectra were acquired in positive and negative ionization mode for pooled samples, collecting a full MS scan from m/z 200-1500 at 70K resolution and ACG target of 3e^6^. The top five most abundant ions per scan were selected for MS/MS with a resolution of 17.5K and ACG target of 1e^5^, and stepped collision energies of 10, 20 and 40 NCE. Maximum ion fill time was 50 ms, dynamic exclusion was 3 sec, and an isolation window of 1 m/z was used.

### Untargeted Data analysis

Positive and negative ionization mode datasets were obtained by acquiring full scan mode data of each sample, as well as data-dependent MS2 data of each pooled sample. Raw data were exported to Compound Discoverer Software (v3.1, Thermo) for deconvolution, alignment and annotation. Putative novel feature dereplication was performed against an in-house database of previously acquired features, using a custom Python script which matched features within a mass and retention time threshold of 5 ppm and 0.2 min, respectively. MS2 data were converted to mzml format using Compound Discoverer and exported to Ometa Labs Flow Analysis Platform. Molecular networking analysis was performed using the Classical Networking workflow (see supplemental information for workflow parameters). MS2 spectra of related compounds were grouped within the dataset according to similarity, and searched against reference spectral libraries (GNPS, NIST, MoNa).

### Extraction Methodology for Orthogonal Fractionation

#### ZYM301

The 8 L of ZYM301 and 4 L of empty plasmid control were processed identically. Bacterial cells were first removed via centrifugation (5000 RPM, 15 min) and discarded. The clarified broth was extracted with a 5% (w/v) addition of activated HP20 resin, and allowed to gently stir overnight. The resin was filtered from the aqueous broth, then extracted with methanol (2 × 1 L), followed by acetone (2 × 1 L). The organic fractions were combined and dried *in vacuo*, yielding a thick aqueous suspension. The aqueous layers were diluted to 500 mL using distilled water, and then partitioned against an equal volume of ethyl acetate four times (4 × 500 mL). The ethyl acetate layer was dried over MgSO_4_, filtered, and finally dried *in vacuo* yielding 522.18 mg of BGC extract, and 249.55 mg of control extract. These extracts were reconstituted in methanol, and partitioned into three roughly equal aliquots for fractionation.

#### ZYM302

ZYM302 (2 L) and the corresponding empty vector control (2 L) were extracted identically. Cells were removed via centrifugation (5000 RPM, 15 min) and discarded. Clarified broth was extracted via liquid-liquid partition using an equal volume of a 4:1 mixture of ethyl acetate and isopropanol (3 × 2 L). Organic layers were combined, and dried *in vacuo* yielding thick brown oily residues for both ZYM302 and its corresponding empty vector control (3.585 g and 1.571 g respectively). This material was partitioned into ~200 mg aliquots for further processing.

### Orthogonal Fractionation

#### Silica fractionation ZYM301

Flash chromatography was performed using a Biotage Selekt automated chromatography system utilizing pre-packed Biotage Sfar HC Duo (10 g) silica columns. Both the empty vector control extract (79.35 mg), and the ZYM301 extract (173.21 mg) were fractionated identically. Material was fractionated using a flow rate of 40 mLmin^−1^, collecting 60 mL fractions (4 CV). Material was eluted using a three-solvent system, consisting of hexanes (solvent A), ethyl acetate (solvent B), and methanol (solvent C). The column was initiated with a linear increasing gradient from 30% to 100% solvent B in solvent A for 12 CVs (F1-F3). This was followed by an isocratic elution using 100% solvent B for 4 CVs (F4). This was followed by another linear increasing gradient from 10% to 80% solvent C in solvent B over 8 CVs (F5-F6). Finally, an 80% isocratic wash of solvent C in solvent B was performed, over 8 CVs. This generated another 2 fractions that were combined into a single final fraction (F7), yielding 7 fractions in total for both extracts. Fractions were dried into pre-weighed vials using a V10-touch evaporator (Biotage) coupled with a Gilson GX-271 Liquid Handler. Fractions were used for both bioactivity assessment and MS-analysis without further purification. For MS-analysis, samples were brought up to a concentration of 1 mgmL^−1^, and 4 μL was injected and run using the UPLC-MS/MS method previously described.

#### Silica fractionation ZYM302

Fractionations were carried out identically for both ZYM302 (204.8 mg) and empty vector control (201.1 mg). Fractionations and MS-analyses were carried out using the same gradient, flow rate, drying procedure, and sample concentrations as previously described for ZYM301. The only difference was 30 mL fractions were collected (2 CV each). Using the same solvents and gradient for elution as described above, the fractions were generated as follows, 12 CVs (F1-F6), 4 CVs (F7-F8), and 8 CVs (F9-F12). The final 8 CVs were divided into 2 × 4 CV blocks (F13-F14), yielding 14 fractions total.

#### C18 Fractionation ZYM301

Flash chromatography was performed using a Biotage Selekt automated chromatography system utilizing pre-packed Biotage Sfar C18 (12 g) columns. Both the empty vector control extract (83.98 mg), and the ZYM301 extract (174.10 mg) were fractionated identically. Material was eluted with a flow rate of 12 mLmin^−1^ collecting 68 mL fractions (4 CV). Material was eluted using a simple 2-solvent gradient system, consisting of H_2_O (solvent A), methanol (solvent B). The column was first washed with 5% methanol in H2O for 4 CV (F1), followed by a linear increasing gradient from 5% to 100% methanol over 20 CV (F2-F6). An isocratic gradient of 100% methanol was then applied for 8 CV, and this wash was combined into one fraction (F7), yielding 7 fractions in total. Fractions were dried into pre-weighed vials using a V10-touch evaporator (Biotage®) coupled with a Gilson GX-271 Liquid Handler. Fractions were used for both bioactivity assessment and MS-analysis without further purification. For MS-analysis, samples were brought up to a concentration of 1 mgmL^−1^, and 4 μL was injected and run using the UPLC-MS/MS method previously described.

#### C18 Fractionation ZYM302

Fractionations were carried out identically for both ZYM302 (208.5 mg) and empty vector control (200.8 mg). Fractionations and MS-analyses were carried out using the same gradient, flow rate, drying procedure, and sample concentrations as ZYM301. The only difference was 34 mL fractions were collected (2 CV each). Using the same solvents and gradient for elution as described above, the fractions were generated as follows, 4 CVs (F1-F2), and 20 CVs (F3-F12). The final 8 CVs were divided into 2 × 4 CV blocks (F13-F14), yielding 14 fractions total.

#### LH20 Size-Exclusion Fractionation ZYM301

Size-exclusion chromatography was performed with a hand packed LH20 column (15.9 × 600 mm). Both the empty vector control extract (86.22 mg), and the ZYM301 extract (174.87 mg) were fractionated identically. Material was eluted using 100% methanol, with a flow rate of ~0.5 mLmin^−1^ collecting 4 mL fractions over 180 mL, yielding 45 initial fractions. Due to mass limitations, fractions 1-10 were combined (F1), and then every 4 fractions from 11-30 (F2-F5), and all remaining fractions (31-45) were combined (F7) yielding 7 fractions in total. Fractions were dried using a Speedvac Vacuum Concentrator, and used without further purification. For MS-analysis, samples were brought up to a concentration of 1 mgmL^−1^, and 4 μL was injected and run using the UPLC-MS/MS method previously described.

### Orthogonal fractionation data analysis

The input features for the PLS analysis were all features from the gene cluster and empty vector sample for which MS2 data was captured. Features that were more than 0.9 cosine similarity were combined into a consensus spectrum. For each feature, the peak area was determined via XIC integration with 0.2 tolerance on the retention time. Peak areas were then normalized to sum to equal amounts for each fraction. For the union of all 38523 features across all 21 fractions (7 fractions for 3 different fractionation methods), the normalized peak areas were cast into a feature matrix of dimension 21 × 38523. The corresponding bioactivity vector of dimension 21 was composed by taking the average inhibition percentage across the three replicates and subtracting the inhibition observed in the same fraction for the empty vector.

A PLS analysis with two components and standard scaling was then run on the resulting feature matrix and bioactivity matrix, and the selectivity ratios were calculated from the resulting PLS vectors. Features with a differential expression ratio (measured by the sum of peak areas across all fractions for the gene cluster sample versus empty vector control) less than 100 were disregarded. The resulting selectivity ratios were tabled and plotted on top of a network plot for all gene cluster and empty vector sample features by scaling the node size for each feature.

The network plot was run with a cosine similarity cutoff of 0.7, 8 matching peaks and a maximum shift of 250.

### Purification

After 7-day culture period, 180L worth culture flasks were combined and centrifuged (4,000 rpm for 10 min). Mycelial portion was discarded. The supernatant was absorbed onto HP20 resin (5%, w/v) for overnight overhead spinning. The resin was filtered and washed with water to remove water soluble components. The resin was extracted in ethyl acetate (12L). The organic phase was dried *in vacuo* to afford 70g of dried crude extract.

Crude extract was subjected to a size exclusion column packed with Sephadex LH-20 and manually fractionated in methanol. The fractions were screened by LC-MS. Fractions containing the compounds of interest were combined and subjected to preparative HPLC purifications. The first round of HPLC was conducted on a C18 column (250 × 10 mm, phenomenex, CH_3_CN-H_2_O, 0.1% FA, flow rate: 8ml/min), and the second round of HPLC was conducted on a phenyl hexyl column (250 × 10 mm, phenomenex, CH_3_CN-H_2_O, 0.1%FA, flow rate: 8ml/min).

HPLC fractions containing the compounds were dried to yield metapeptin A (25mg) and metapeptin B (1.8mg).

Metapeptin A, white, amorphous powder, (+)-HR-ESIMS m/z = 867.5224, [M + H]^+^ (calcd for C_46_H_71_N_6_O_10_^+^, 867.5226)

Metapeptin B, white, amorphous powder, (+)-HR-ESIMS m/z = 853.5069, [M + H]^+^ (calcd for C_45_H_69_N_6_O_10_^+^, 853.5070)

### NMR

All NMR spectra were recorded on a Bruker AVANCE at 900 MHz (^1^H NMR) and 226 MHz (^13^C NMR). NMR spectra were analyzed using Mestrenova 9.0.1.

### Monomer synthesis

The NMe-Monomer was designed with all natural (L) stereochemistry based upon bioinformatic prediction from the encoded BGC analysis. The molecule was synthesized in 8 steps as illustrated in Figure S6. Detailed synthetic procedures and characterization are available upon request.

### Molecular modeling

Most simulations were carried out with Schrodinger’s Small Molecule Drug Discovery Suite version 2022-1. Glide docking was done with the XP precision level with post-docking minimization and applying strain correction terms. MM-GBSA calculations used the VSGB solvation model with OPLS4 force field. Induced-fit docking was completed with the Extended Sampling protocol generating up to 80 poses, residues within 15 Å of the binding site were refined, and the docking was done with SP precision. Jaguar was used for the pKa calculations using B3LYP-D3 level of theory and the 6-31G** basis set. Maestro was used for computing dipoles and making the docking poses picture. The dipole result was a property calculated by QikProp. Desmond was used for the molecular dynamics simulations using the NPT ensemble class at a temperature of 300 K for 25ns. PDB ID 6LZC was used for the HsMetAP1 protein.

## References

1. Behan, F. M. et al. Prioritization of cancer therapeutic targets using CRISPR–Cas9 screens. Nature 568, 511–516 (2019).

2. Hahn, W. C. et al. An expanded universe of cancer targets. Cell 184, 1142–1155 (2021).

3. Brown, D. G. & Wobst, H. J. A Decade of FDA-Approved Drugs (2010-2019): Trends and Future Directions. J. Med. Chem. 64, 2312–2338 (2021).

4. Stone, S., Newman, D. J., Colletti, S. L. & Tan, D. S. Cheminformatic analysis of natural product-based drugs and chemical probes. Nat. Prod. Rep. 39, 20–32 (2022).

5. Newman, D. J. & Cragg, G. M. Natural Products as Sources of New Drugs over the Nearly Four Decades from 01/1981 to 09/2019. J. Nat. Prod. 83, 770–803 (2020).

6. Bergner, A. et al. KRAS Binders Hidden in Nature. Chem. Weinh. Bergstr. Ger. 25, 12037–12041 (2019).

7. Harvey, A. L., Edrada-Ebel, R. & Quinn, R. J. The re-emergence of natural products for drug discovery in the genomics era. Nat. Rev. Drug Discov. 14, 111–129 (2015).

8. Li, J., Amatuni, A. & Renata, H. Recent advances in the chemoenzymatic synthesis of bioactive natural products. Curr. Opin. Chem. Biol. 55, 111–118 (2020).

9. Lipinski, C. A. Rule of five in 2015 and beyond: Target and ligand structural limitations, ligand chemistry structure and drug discovery project decisions. Adv. Drug Deliv. Rev. 101, 34–41 (2016).

10. Chevrette, M. G. et al. Evolutionary dynamics of natural product biosynthesis in bacteria. Nat. Prod. Rep. 37, 566–599 (2020).

11. Atanasov, A. G., Zotchev, S. B., Dirsch, V. M., the International Natural Product Sciences Taskforce & Supuran, C. T. Natural products in drug discovery: advances and opportunities. Nat. Rev. Drug Discov. 20, 200–216 (2021).

12. Henrich, C. J. & Beutler, J. A. Matching the power of high throughput screening to the chemical diversity of natural products. Nat. Prod. Rep. 30, 1284–1298 (2013).

13. Rappé, M. S. & Giovannoni, S. J. The uncultured microbial majority. Annu. Rev. Microbiol. 57, 369–394 (2003).

14. Handelsman, J., Rondon, M. R., Brady, S. F., Clardy, J. & Goodman, R. M. Molecular biological access to the chemistry of unknown soil microbes: a new frontier for natural products. Chem. Biol. 5, R245–249 (1998).

15. Katz, M., Hover, B. M. & Brady, S. F. Culture-independent discovery of natural products from soil metagenomes. J. Ind. Microbiol. Biotechnol. 43, 129–141 (2016).

16. Stevenson, L. J., Owen, J. G. & Ackerley, D. F. Metagenome Driven Discovery of Nonribosomal Peptides. ACS Chem. Biol. 14, 2115–2126 (2019).

17. Stevenson, L. J. et al. Metathramycin, a new bioactive aureolic acid discovered by *heterologous expression of a metagenome derived biosynthetic pathway*. RSC Chem. Biol. 2, 556–567 (2021).

18. Peek, J. et al. Rifamycin congeners kanglemycins are active against rifampicin-resistant bacteria via a distinct mechanism. Nat. Commun. 9, 4147 (2018).

19. Owen, J. G. et al. Multiplexed metagenome mining using short DNA sequence tags *facilitates targeted discovery of epoxyketone proteasome inhibitors*. Proc. Natl. Acad. Sci. U. S. A. 112, 4221–4226 (2015).

20. Chang, F.-Y., Ternei, M. A., Calle, P. Y. & Brady, S. F. Discovery and synthetic refactoring of tryptophan dimer gene clusters from the environment. J. Am. Chem. Soc. 135, 17906–17912 (2013).

21. Wang, Z., Forelli, N., Hernandez, Y., Ternei, M. & Brady, S. F. Lapcin, a potent dual topoisomerase I/II inhibitor discovered by soil metagenome guided total chemical synthesis. Nat. Commun. 13, 842 (2022).

22. Xu, G. et al. Combined assembly of long and short sequencing reads improve the efficiency of exploring the soil metagenome. BMC Genomics 23, 37 (2022).

23. Delahaye, C. & Nicolas, J. Sequencing DNA with nanopores: Troubles and biases. PloS One 16, e0257521 (2021).

24. Tedersoo, L., Albertsen, M., Anslan, S. & Callahan, B. Perspectives and Benefits of High-Throughput Long-Read Sequencing in Microbial Ecology. Appl. Environ. Microbiol. 87, e0062621 (2021).

25. Frottin, F. et al. MetAP1 and MetAP2 drive cell selectivity for a potent anti-cancer agent in synergy, by controlling glutathione redox state. Oncotarget 7, 63306–63323 (2016).

26. Chen, I.-M. A. et al. The IMG/M data management and analysis system v.6.0: new tools and advanced capabilities. Nucleic Acids Res. 49, D751–D763 (2021).

27. Blin, K. et al. antiSMASH 5.0: updates to the secondary metabolite genome mining pipeline. Nucleic Acids Res. 47, W81–W87 (2019).

28. Gavriilidou, A. et al. Compendium of specialized metabolite biosynthetic diversity encoded in bacterial genomes. Nat. Microbiol. 7, 726–735 (2022).

29. Yan, Y., Liu, N. & Tang, Y. Recent developments in self-resistance gene directed natural product discovery. Nat. Prod. Rep. 37, 879–892 (2020).

30. Kale, A. J., McGlinchey, R. P., Lechner, A. & Moore, B. S. Bacterial self-resistance to the natural proteasome inhibitor salinosporamide A. ACS Chem. Biol. 6, 1257–1264 (2011).

31. Culp, E. J. et al. ClpP inhibitors are produced by a widespread family of bacterial gene clusters. Nat. Microbiol. 7, 451–462 (2022).

32. Wenzel, S. C. et al. Production of the Bengamide Class of Marine Natural Products in *Myxobacteria: Biosynthesis and Structure-Activity Relationships*. Angew. Chem. Int. Ed Engl. 54, 15560–15564 (2015).

33. Kautsar, S. A. et al. MIBiG 2.0: a repository for biosynthetic gene clusters of known function. Nucleic Acids Res. gkz882 (2019) doi:10.1093/nar/gkz882.

34. Caesar, L. K., Kellogg, J. J., Kvalheim, O. M. & Cech, N. B. Opportunities and Limitations for Untargeted Mass Spectrometry Metabolomics to Identify Biologically Active Constituents in Complex Natural Product Mixtures. J. Nat. Prod. 82, 469–484 (2019).

35. Nothias, L.-F. et al. Bioactivity-Based Molecular Networking for the Discovery of Drug Leads in Natural Product Bioassay-Guided Fractionation. J. Nat. Prod. 81, 758–767 (2018).

36. Kellogg, J. J. et al. Biochemometrics for Natural Products Research: Comparison of Data Analysis Approaches and Application to Identification of Bioactive Compounds. J. Nat. Prod. 79, 376–386 (2016).

37. Fukuda, T. et al. New Beauvericins, Potentiators of Antifungal Miconazole Activity, Produced by Beauveria sp. FKI-1366: I. Taxonomy, Fermentation, Isolation and Biological Properties. J. Antibiot. (Tokyo) 57, 110–116 (2004).

38. White, K. N., Tenney, K. & Crews, P. The Bengamides: A Mini-Review of Natural Sources, Analogues, Biological Properties, Biosynthetic Origins, and Future Prospects. J. Nat. Prod. 80, 740–755 (2017).

39. Lloyd, K. G., Steen, A. D., Ladau, J., Yin, J. & Crosby, L. Phylogenetically Novel Uncultured Microbial Cells Dominate Earth Microbiomes. mSystems 3, e00055–18 (2018).

40. Brady, S. F. Construction of soil environmental DNA cosmid libraries and screening for clones that produce biologically active small molecules. Nat. Protoc. 2, 1297–1305 (2007).

41. Nurk, S., Meleshko, D., Korobeynikov, A. & Pevzner, P. A. metaSPAdes: a new versatile metagenomic assembler. Genome Res. 27, 824–834 (2017).

42. Hyatt, D. et al. Prodigal: prokaryotic gene recognition and translation initiation site identification. BMC Bioinformatics 11, 119 (2010).

43. Menzel, P., Ng, K. L. & Krogh, A. Fast and sensitive taxonomic classification for metagenomics with Kaiju. Nat. Commun. 7, 11257 (2016).

44. Eddy, S. R. Profile hidden Markov models. Bioinforma. Oxf. Engl. 14, 755–763 (1998).

45. Mistry, J. et al. Pfam: The protein families database in 2021. Nucleic Acids Res. 49, D412–D419 (2021).

46. Steinegger, M. & Söding, J. MMseqs2 enables sensitive protein sequence searching for the analysis of massive data sets. Nat. Biotechnol. 35, 1026–1028 (2017).

47. Navarro-Muñoz, J. C. et al. A computational framework to explore large-scale biosynthetic diversity. Nat. Chem. Biol. 16, 60–68 (2020).

48. Gietz, R. D. & Schiestl, R. H. High-efficiency yeast transformation using the LiAc/SS carrier DNA/PEG method. Nat. Protoc. 2, 31–34 (2007).

