## Supplementary Figures, Tables, and Text for "Precision Discovery of Novel Inhibitors of Cancer Target HsMetAP1 from Vast Metagenomic Diversity"

**This PDF file includes:**

Figs. S1 to S7
Tables S1 to S5
Supplementary Text

FIGURE S1

A

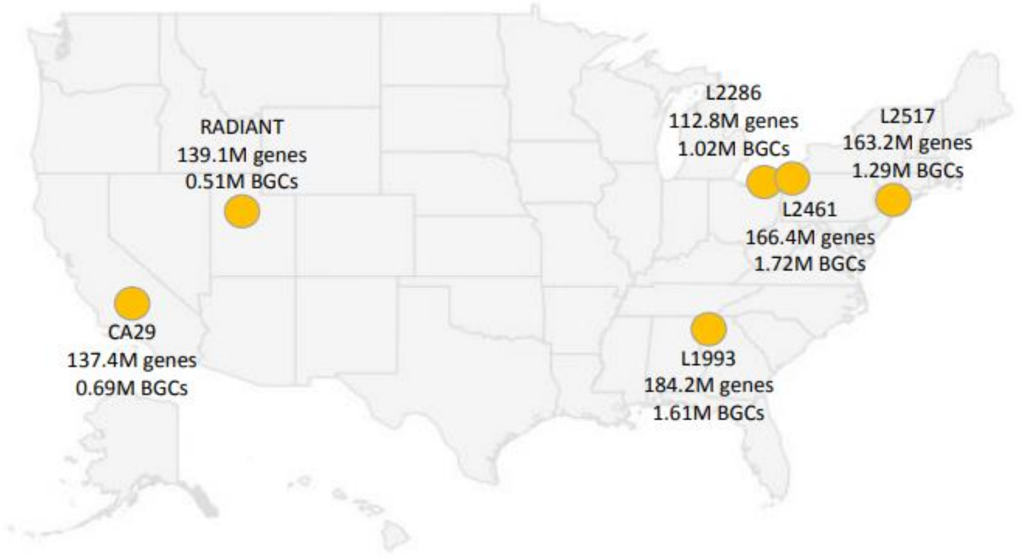


B

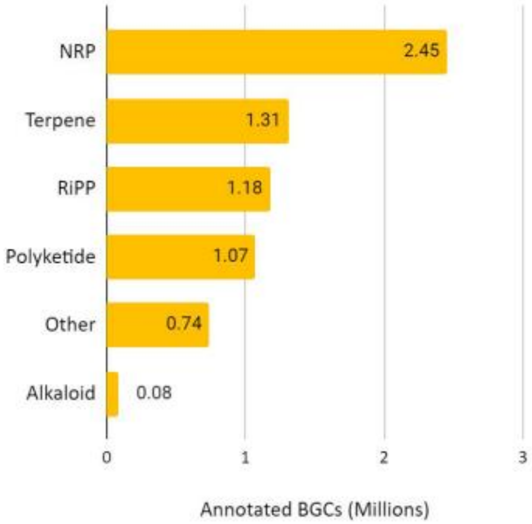


**Figure S1**. **Metagenomic libraries come from diverse locations and contain rich biosynthetic potential.** **(A)** Geographic distribution of the 6 soil metagenomic libraries described in this paper. The number of genes represents clusters at 90% amino acid identity. (**B)** NRP, terpene, RiPP, and polyketide clusters are the most common major biosynthetic gene classes represented in the metagenomic libraries

FIGURE S2

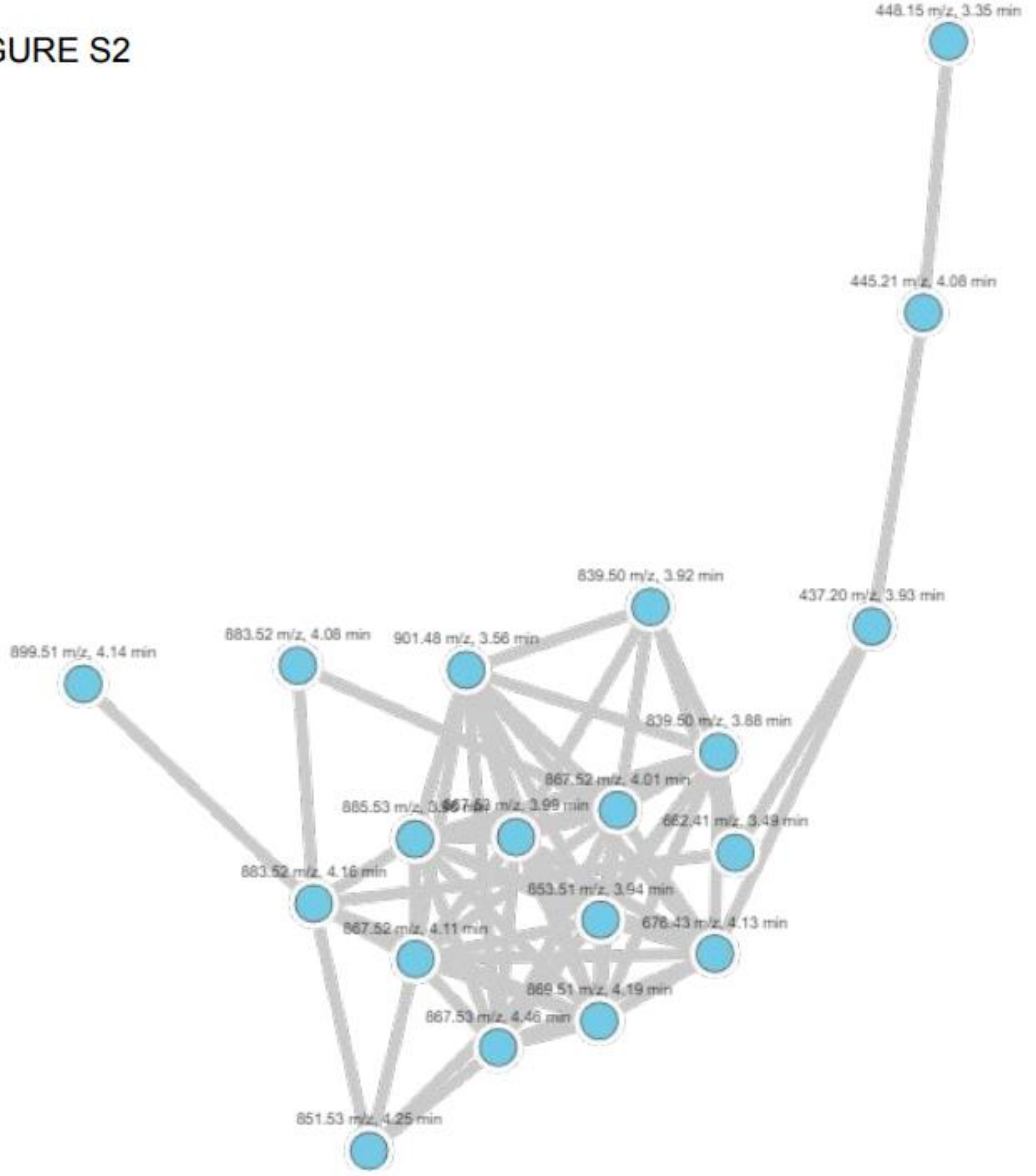


**Figure S2**. **Molecular network of putative novel features expressed in *S.albus:ZYM301* strain.**

FIGURE S3

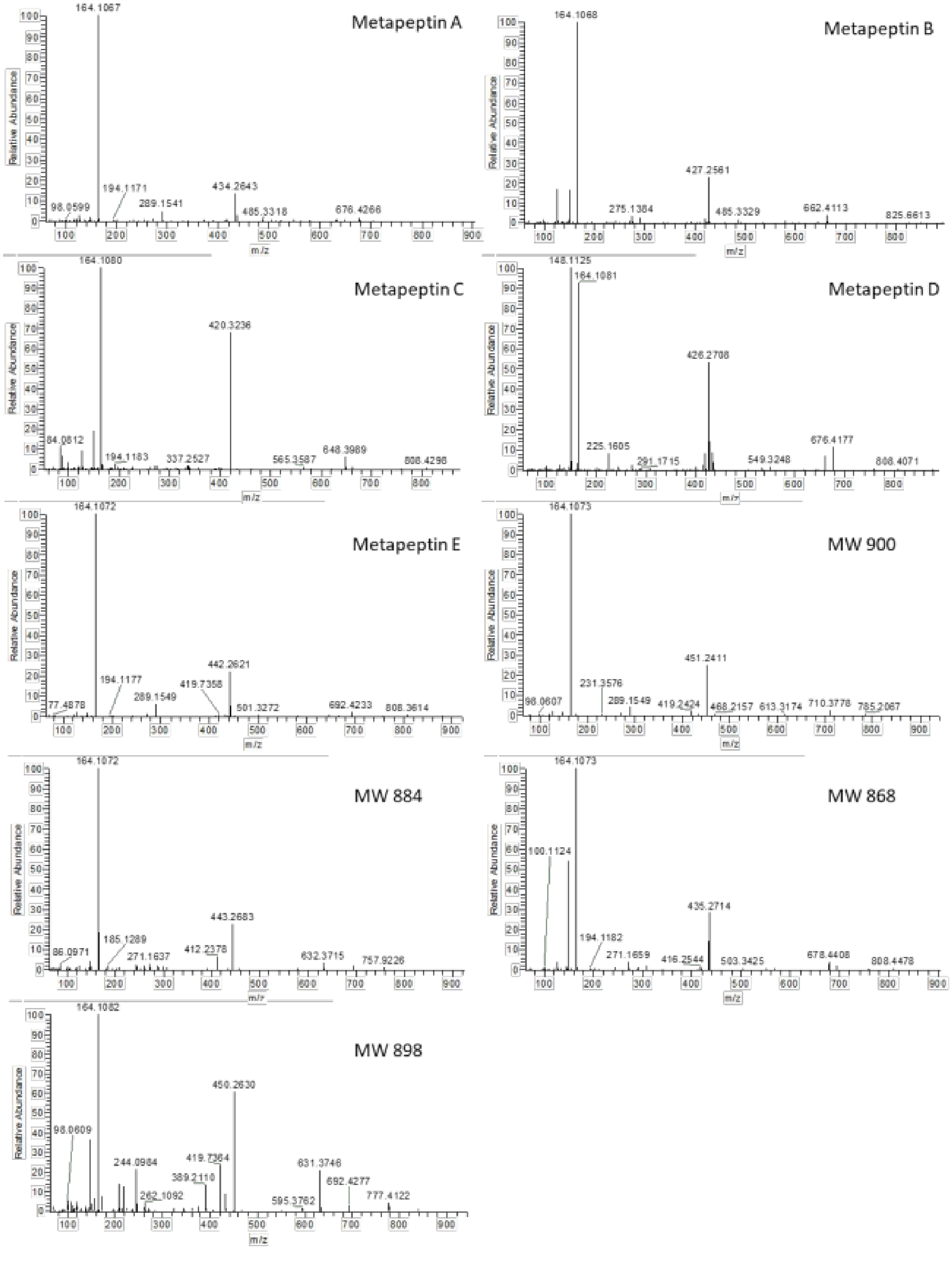


**Figure S3. MS2 spectra for nine of the novel compounds detected in *S.albus:ZYM301*.** All molecules have a common MS2 fragment of m/a 164.1082, suggesting an N-methylated tyrosine residue.

FIGURE S4

A

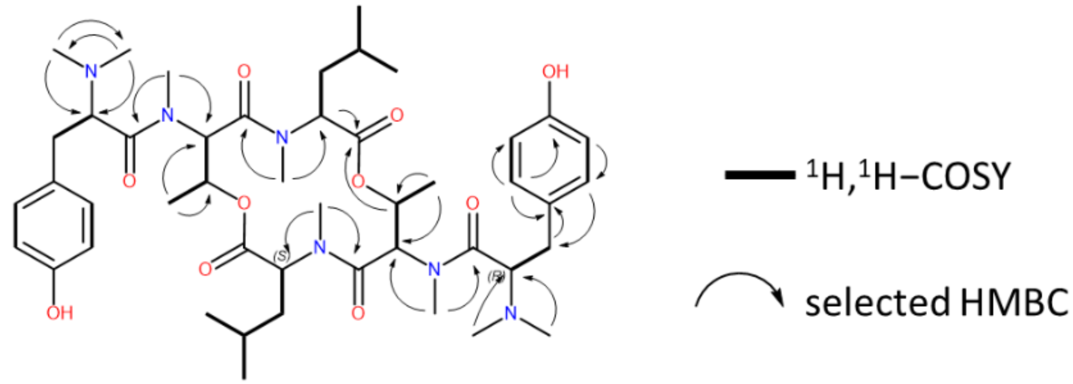

**FIGURE S4 (continued)**

**B**

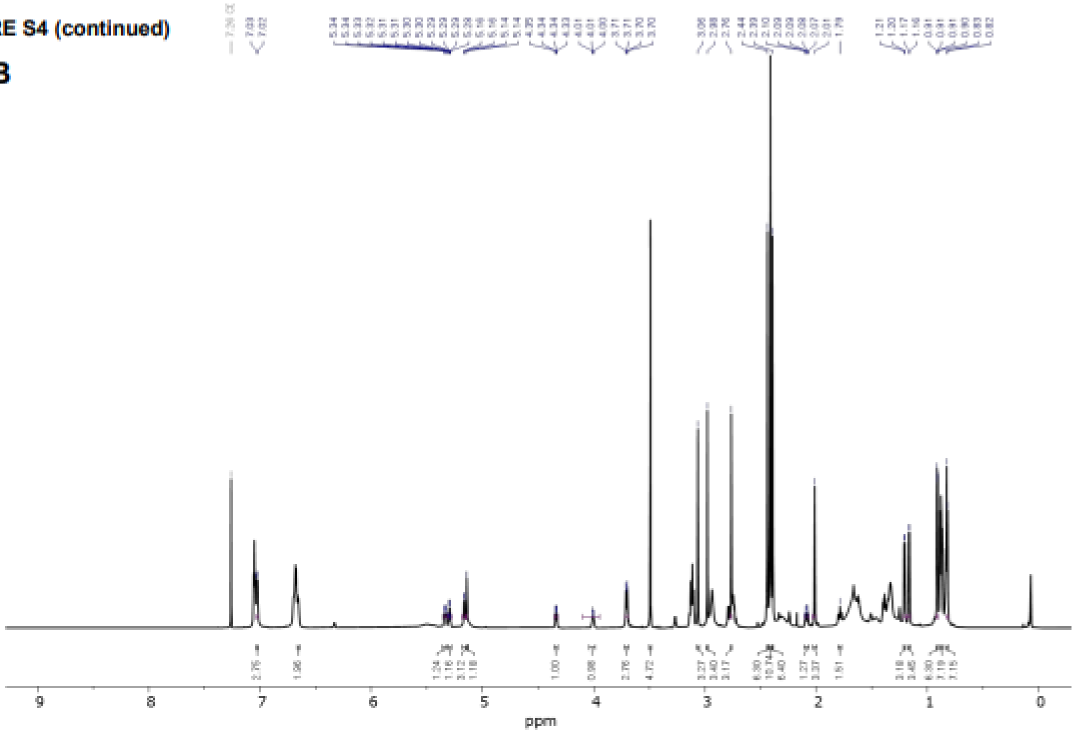


$^{1}$H NMR spectrum

**C**

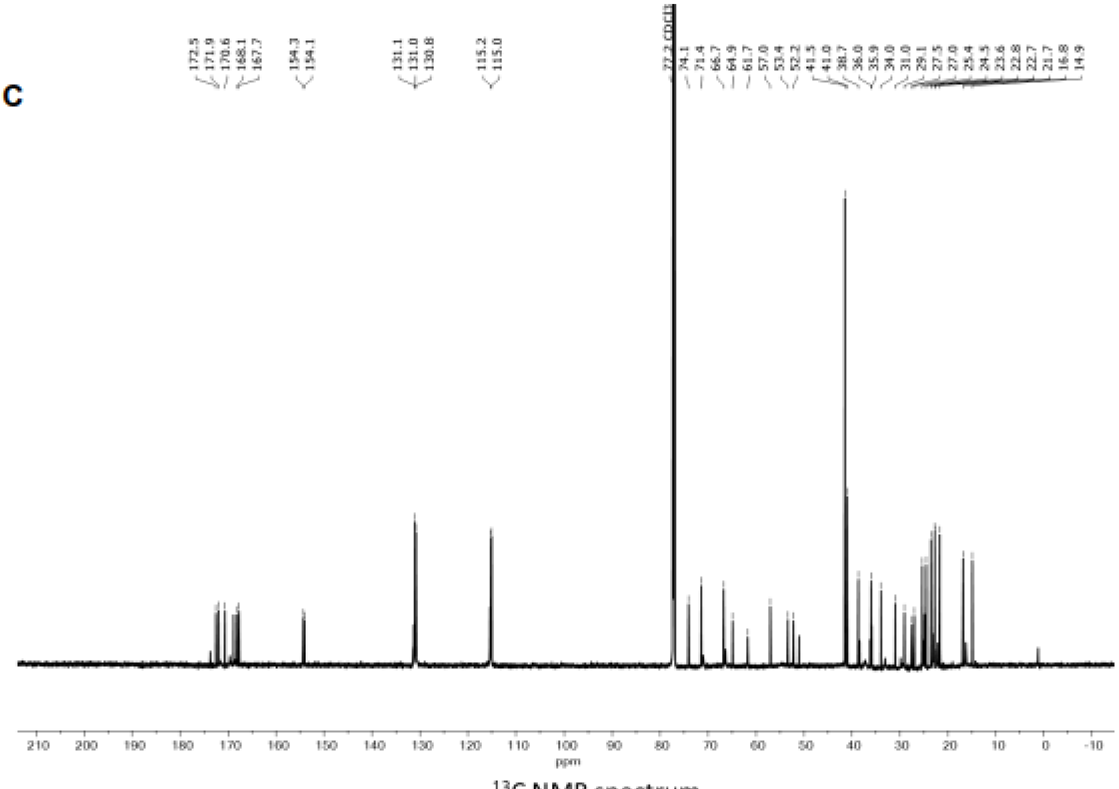


$^{13}$C NMR spectrum

FIGURE S4 (continued)

D

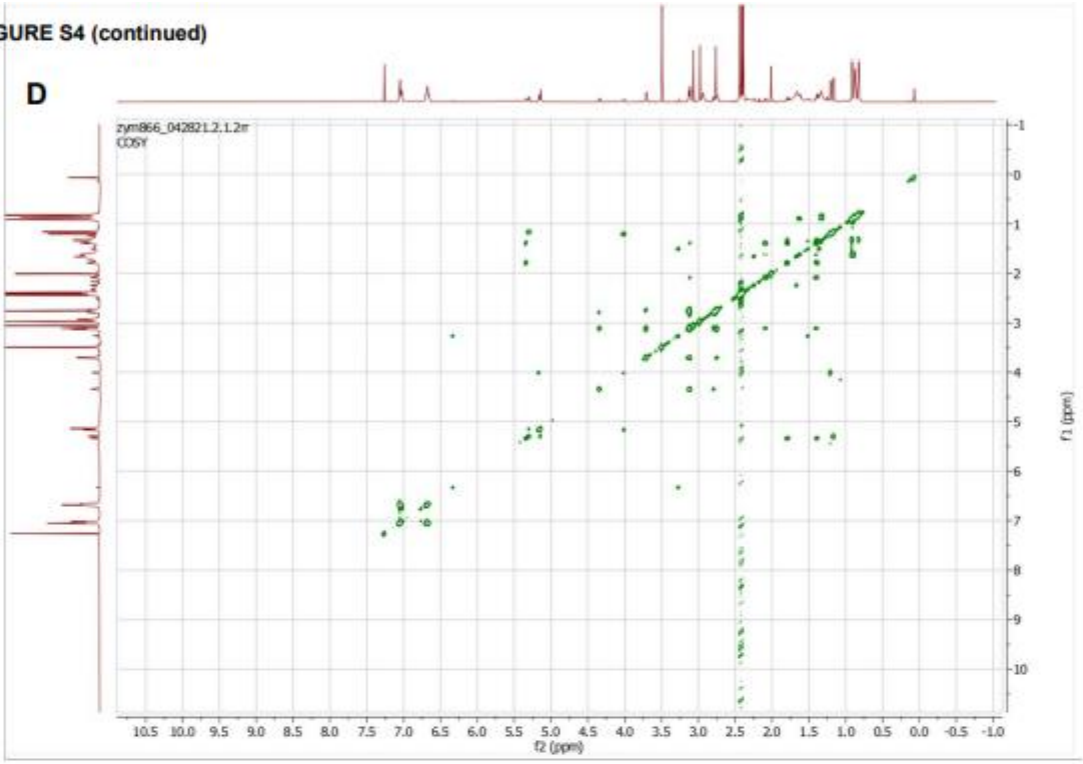


$^1H$–$^1H$ COSY spectrum

E

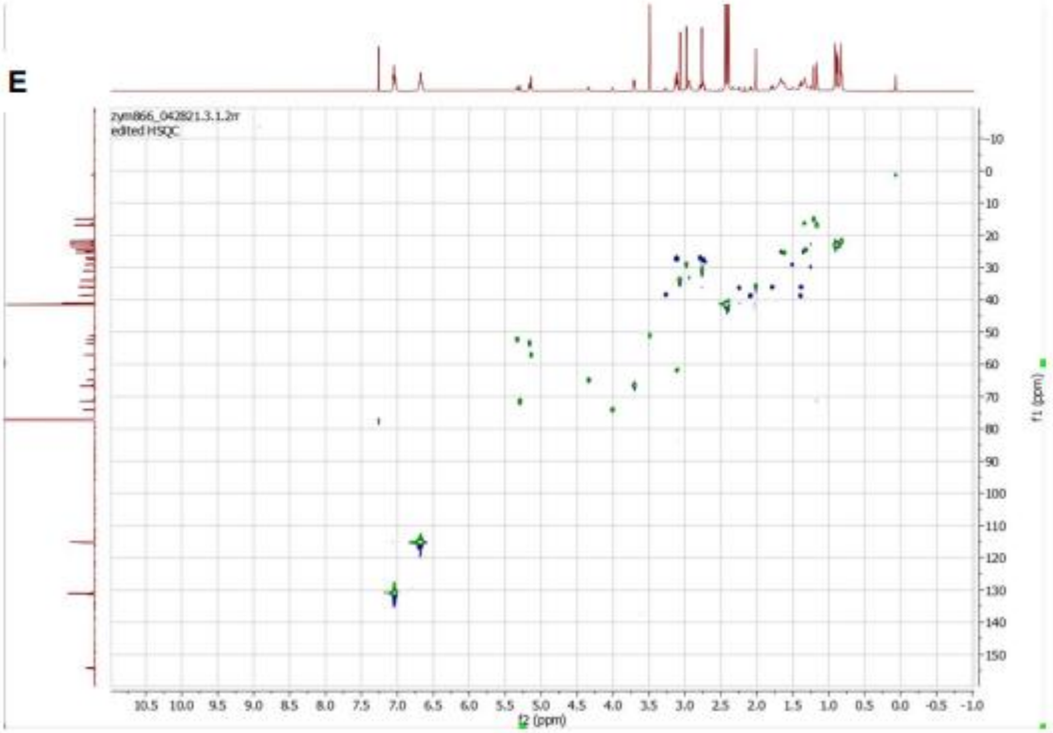


$^1H$– $^{13}C$ HSQC spectrum

**FIGURE S4 (continued)**

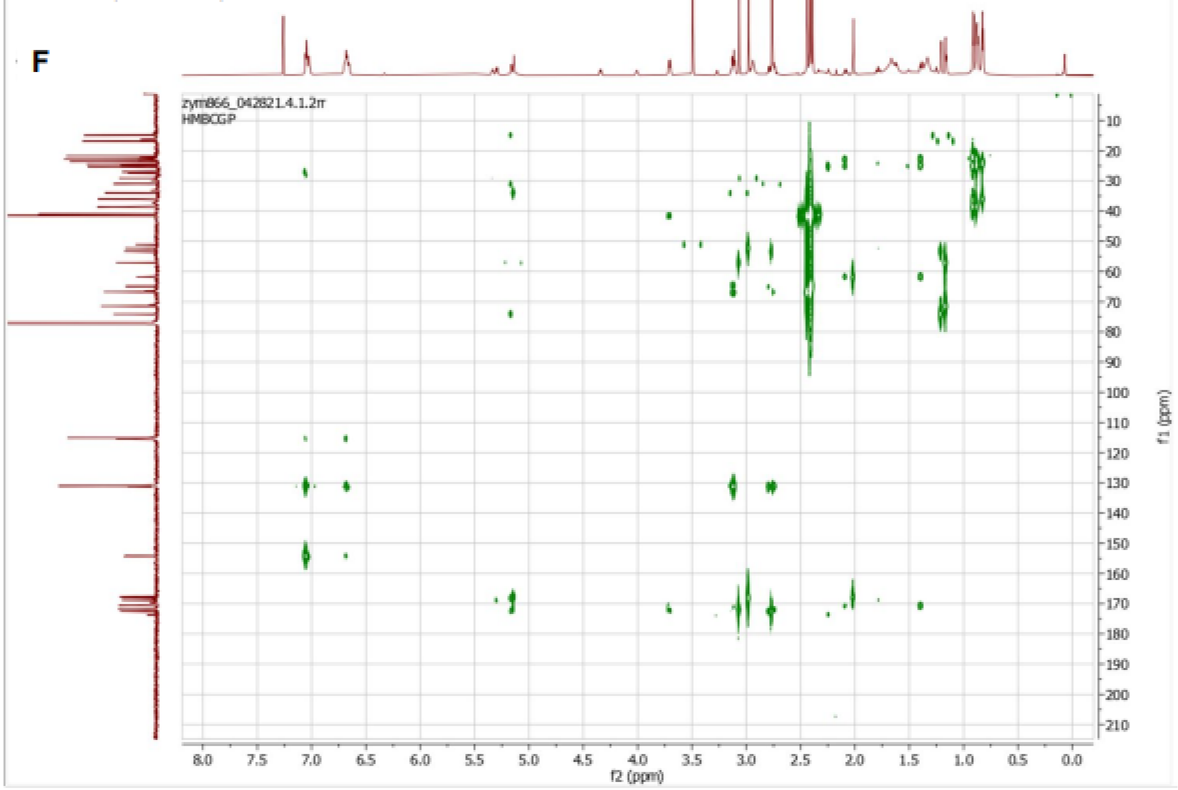


$^{1}H–^{13}C$ HMBC spectrum

**Figure S4. Structure elucidation of metapeptin A**. 1D and 2D NMR spectra of metapeptin A. (**A)** NMR-based key correlations for structural assignment. **(B)** 1H NMR spectrum. **(C)** 13C NMR spectrum. **(D)** 1H-1H COSY spectrum. **(E)** 1H-13C HSQC spectrum. **(F)** 1H-13C HMBC spectrum. Recording with 900 MHz for 1H and 226 MHz for 13C, cryo probe, in CDCl3

FIGURE S5

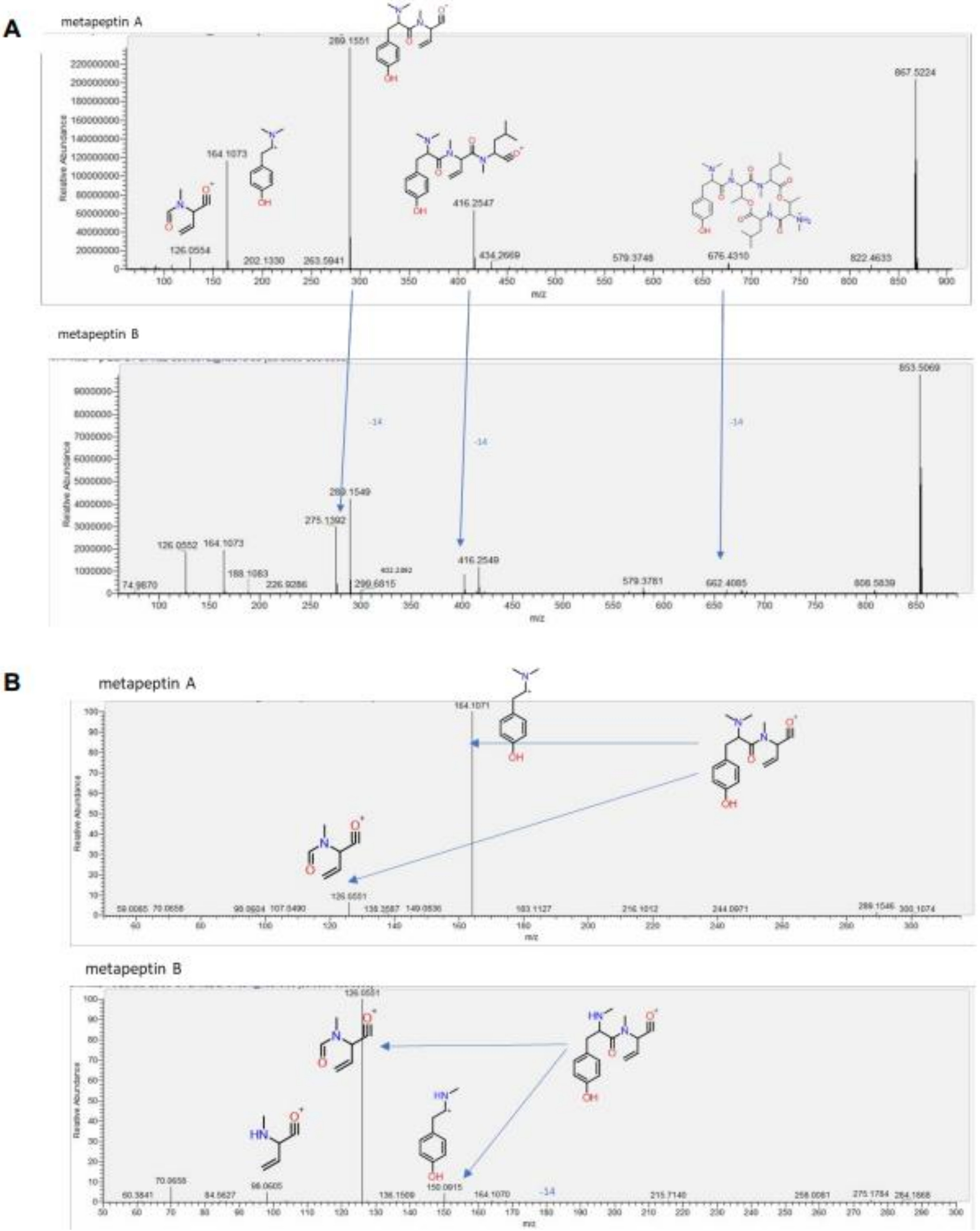


**Figure S5**. **Structure elucidation of metapeptin B. (A)**, HR-MS/MS fragmentation pattern of metapeptin A (upper) and metapeptin B (lower). **(B)** HR-MS/MS fragmentation pattern of N,N-dimethyl-Try-N-methyl-Thr moiety of metapeptin A (upper) and N-methyl-Try-N-methyl-Thr moiety of metapeptin B (lower). **C)**, the major fragmentations species from HR-MS/MS measurement of metapeptin A. **D)**, the major fragmentations species from HR-MS/MS measurement of metapeptin B.

FIGURE S6

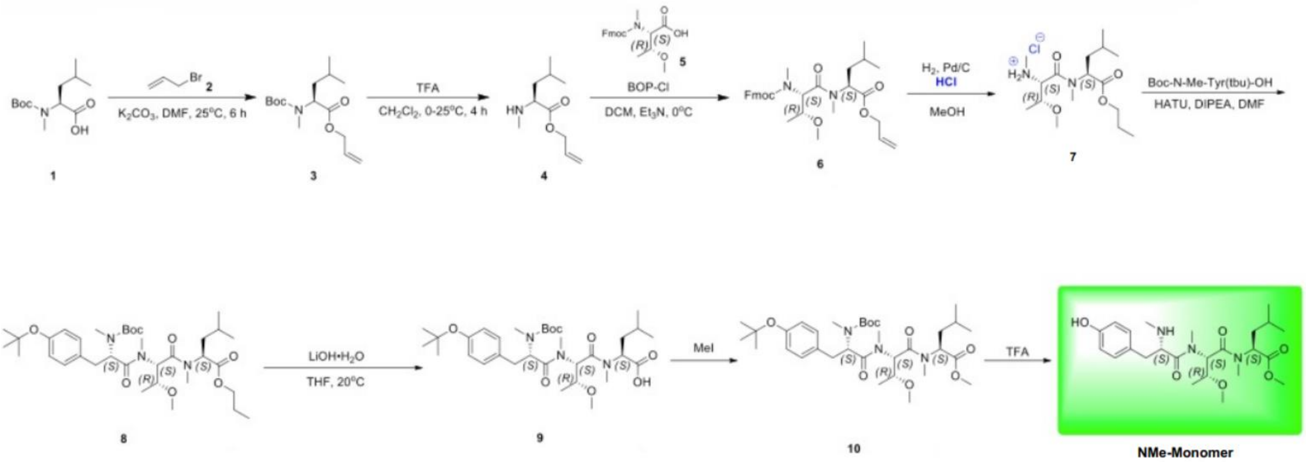


**Figure S6. Synthesis scheme for the NMe-Monomer.** The NMe-Monomer was designed with all natural (L) stereochemistry based upon bioinformatic prediction from the encoded BGC analysis

FIGURE S7

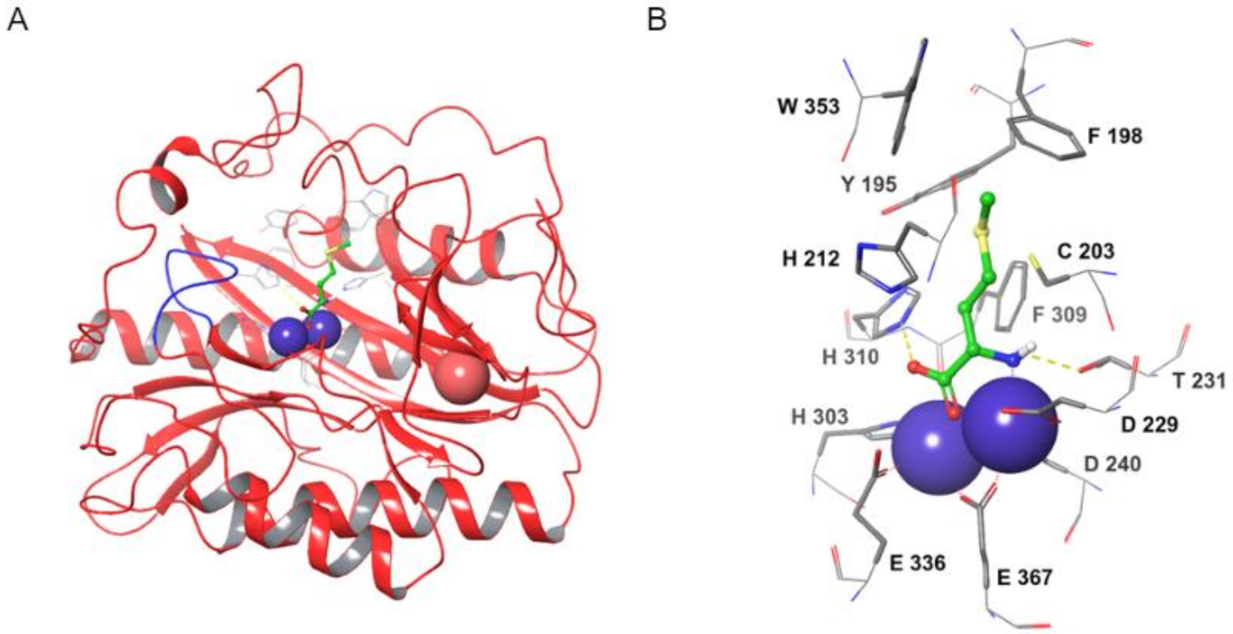


**Figure S7. (A)** Crystal structure of *Hs*MetAP1 (PDB ID 4u6j) in complex with a methionine substrate (green carbon atoms, stick representation). The catalytic domain adopts a pita-bread. The loop (blue) is in the active conformation opening the active site. **(B)** The active site has two divalent ions complex with four negatively charged amino acids (D229, D240, E336, and E367) . The methionine substrate is shown with green caron atoms in sticks representation.

#### TABLE S1

| Cluster | Phylum | Class | Cluster size (bp) |
|---|---|---|---|
| ZYM301 | Actinobacteria | nrps | 34,196 |
| ZYM302 | Actinobacteria | nrps-like | 50,832 |
| ZYM303 | Ignavibacteriae | nrps | 29,675+ |
| ZYM304 | Actinobacteria | nrps | 22,859+ |
| ZYM305 | Actinobacteria | nrps | 2,910 |
| ZYM306 | Actinobacteria | nrps | 4,813+ |
| ZYM307 | Actinobacteria | betalactone | 28,828 |
| ZYM308 | Zixibacteria | nrps | 24,522+ |
| ZYM309 | Acidobacteria | ladderane,arylpolyene | 28,788+ |
| ZYM310 | Unknown | nrps,t1pks | 30,842+ |
| ZYM311 | Actinobacteria | betalactone | 27,623 |
| ZYM312 | Actinobacteria | nrps | 32,336+ |
| ZYM313 | Actinobacteria | nrps,indole | 148,500 |
| ZYM314 | Actinobacteria | nrps | 37,733 |
| ZYM315 | Proteobacteria | t1pks,nrps | 59,910 |
| ZYM316 | Actinobacteria | betalactone | 22,931+ |
| ZYM317 | Bacteroidota | nrps | 31,533+ |
| ZYM318 | Actinobacteria | nrps,betalactone | 31,963 |
| ZYM319 | Actinobacteria | nrps,t1pks | 26,724 |
| ZYM320 | Actinobacteria | nrps,terpene | 87,330 |
| ZYM321 | Actinobacteria | pks-like | 31,352+ |
| ZYM322 | Actinobacteria | nrps-like,pks-like,betalactone,terpene | 27,666 |
| ZYM323 | Actinobacteria | nrps | 73,939+ |
| ZYM324 | Proteobacteria | ladderane | 3,348+ |
| ZYM325 | Actinobacteria | betalactone | 27,775 |
| ZYM326 | Actinobacteria | nrps-like | 21,982 |
| ZYM327 | Actinobacteria | pks-like | 33,473+ |
| ZYM328 | Actinobacteria | nrps | 34,298 |
| ZYM329 | Proteobacteria | betalactone | 24,920 |
| ZYM330 | Actinobacteria | nrps | 23,230+ |
| ZYM331 | Actinobacteria | nrps | 56,073 |
| ZYM332 | Actinobacteria | nrps | 8,102+ |
| ZYM333 | Proteobacteria | nrps | 28,499+ |
| ZYM334 | Actinobacteria | terpene,nrps | 87,329 |
| ZYM335 | Actinobacteria | t2pks | 41,587 |

**TABLE S1. Clusters found with a MetAP resistance gene**. Phylum was assessed by Kaiju (ref), class was assigned by antiSMASH, and the size of the cluster is the ‘region’ identified by antiSMASH

#### TABLE S2

| ORF | Residues | Proposed Function | Closest relative | Identity (%) | Accession |
|---|---|---|---|---|---|
| *orf1* | 227 | Unknown | hypothetical protein [Mycobacterium sp. NAZ190054] | 61.47 | WP_067958064.1 |
| *orf2* | 176 | Transcriptional Regulator | MarR family transcriptional regulator [Stackebrandtia nassauensis] | 72.05 | WP_013018374.1 |
| *orf3* | 292 | Peptidase | M56 family metallopeptidase [Actinophytocola xanthii] | 67.00 | WP_075127515.1 |
| *orf4* | 131 | Transcriptional Regulator | CopY family transcriptional regulator [Actinophytocola xanthii] | 83.61 | OLF15564.1 |
| *orf5* | 377 | Transporter | MFS transporter [Actinophytocola xanthii] | 78.18 | WP_075127531 |
| *orf6* | 230 | Regulator | GntR family transcriptional regulator [Nonomuraea guangzhouensis] | 73.25 | WP_219538193.1 |
| *orf7* | 368 | Transporter | cation diffusion facilitator family transporter [Saccharopolyspora sp. ASAGF58] | 83.29 | WP_168588145.1 |
| *mtpA* | 390 | Phosphoesterase | metallophosphoesterase [Actinophytocola algeriensis] | 79.37 | WP_184811708.1 |
| *mtpB* | 3978 | Non-ribosomal peptide synthetase | non-ribosomal peptide synthetase [Streptomyces sp. 769] | 51.75 | WP_039638096.1 |
| *mtpC* | 546 | Transporter | ABC transporter substrate-binding protein [Actinophytocola oryzae] | 74.82 | WP_133906182.1 |
| *mtpD* | 252 | Methyltransferase | class I SAM-dependent methyltransferase [uncultured bacterium] | 85.26 | AXL05822.1 |
| *mtpE* | 78 | Agmatinase | agmatinase family protein [Lentzea indica] | 81.71 | WP_167978039.1 |
| *mtpF* | 269 | Methionine aminopeptidase | type I methionyl aminopeptidase [Saccharopolyspora erythraea] | 92.02 | WP_211899443.1 |
| *orf8* | 88 | Transcriptional Regulator | helix-turn-helix domain-containing protein [Kibdelosporangium banguiense] | 92.05 | WP_209643757.1 |
| *orf9* | 155 | Unknown | hypothetical protein SAMN04488564_112257 [Lentzea waywayandensis] | 74.19 | SFR28146.1 |
| *orf10* | 140 | Transcriptional Regulator | TetR/AcrR family transcriptional regulator [Amycolatopsis mediterranei] | 87.50 | WP_013227486.1 |

**TABLE S2. Proposed functions of the genes in the ZYM301 (metapeptin) gene cluster and their closest relatives.**

### TABLE S3

| ORF | Residues | Proposed Function | Closest relative | Identity (%) | Accession |
|---|---|---|---|---|---|
| *orf1* | 88 | Methyltransferase | SAM-dependent methyltransferase [Prauserella shujinwangii] | 54.55 | WP_245901162 |
| *orf2* | 321 | Prenyltransferase | UbiA family prenyltransferase [Amycolatopsis anabasis] | 49.15 | WP_246257859.1 |
| *orf3* | 414 | Oxidase | cytochrome P450 [Qaidamihabitans albus] | 59.66 | WP_199430517.1 |
| *orf4* | 547 | Prenyltransferase | prenyltransferase [Amycolatopsis anabasis] | 61.10 | WP_158888231.1 |
| *orf5* | 277 | Methyltransferase | SAM-dependent methyltransferase [Kutzneria sp. CA-103260] | 62.83 | WP_211768627.1 |
| *orf6* | 141 | Unknown | hypothetical protein [Actinophytocola xanthii] | 72.18 | WP_075128003.1 |
| *orf7* | 289 | Carboxylesterase | pimeloyl-ACP methyl ester carboxylesterase [Saccharopolyspora phatthalungensis] | 80.56 | MBB5156950.1 |
| *orf8* | 1723 | 6-MSA synthase | acyltransferase domain-containing protein [Nocardia sp. ncl2] | 84.94 | WP_216893689.1 |
| *orf9* | 344 | Ketoacyl-ACP synthase | ketoacyl-ACP synthase III family protein [Nocardia sp. ncl2] | 80.42 | WP_216893690.1 |
| *orf10* | 378 | Unknown | DUF1205 domain-containing protein [Nocardia albiluteola] | 89.28 | WP_215917067.1 |
| *orf11* | 57 | Unknown | No significant similarity | N/A | N/A |
| *orf12* | 195 | Regulator | TetR/AcrR family transcriptional regulator [Solihabitans fulvus] | 83.59 | WP_149852387.1 |
| *orf13* | 279 | Methyltransferase | class I SAM-dependent methyltransferase [Solihabitans fulvus] | 88.55 | WP_149852430.1 |
| *orf14* | 103 | Unknown | pilus assembly protein TadE [Actinophytocola sp.] | 81.40 | MPZ79671.1 |
| *orf15* | 495 | Oxidase | copper oxidase [Actinophytocola xanthii] | 89.46 | OLF14973.1 |
| *orf16* | 141 | Unknown | SRPBCC family protein [Actinophytocola xanthii] | 75.41 | WP_075128001.1 |
| *orf17* | 258 | Unknown | hypothetical protein AVL59_18415 [Streptomyces griseochromogenes] | 45.11 | ANP51334.1 |
| *orf18* | 944 | Regulator | AAA family ATPase [Actinophytocola xanthii] | 84.23 | WP_075128000.1 |
| *orf19* | 414 | MDR | medium chain dehydrogenase/reductase family protein [Streptomyces sp. GMY02] | 78.84 | WP_217307496.1 |
| *orf20* | 222 | Methyltransferase | class I SAM-dependent methyltransferase [Saccharopolyspora erythraea] | 94.59 | QUH02008.1 |
| *orf21* | 93 | Regulator | helix-turn-helix domain-containing protein [Saccharopolyspora sp. K220] | 94.32 | WP_242191135.1 |
| *orf22* | 264 | Methionine aminopeptidase | type I methionyl aminopeptidase [Saccharopolyspora erythraea] | 94.70 | WP_211899443.1 |

**TABLE S3 (continued).**

| ORF | Residues | Proposed Function | Closest relative | Identity (%) | Accession |
|---|---|---|---|---|---|
| *orf23* | 397 | Aminotransferase | aminotransferase class I/II-fold pyridoxal phosphate-dependent enzyme [Vulgatibacter incomptus] | 62.53 | WP_050726861.1 |
| *orf24* | 301 | Oxidoreductase | NAD(P)/FAD-dependent oxidoreductase [Labilithrix sp.] | 63.51 | MBX3188560.1 |
| *orf25* | 746 | NRPS-like | amino acid adenylation domain-containing protein [Moritella yayanosii] | 49.28 | WP_112713926.1 |
| *orf26* | 334 | Oxidoreductase | isopenicillin N synthase family oxygenase [Moritella yayanosii] | 60.37 | WP_112713920.1 |
| *orf27* | 337 | Methyltransferase | methyltransferase [Longimycelium tulufanense] | 50.76 | GGM71615.1 |
| *orf28* | 91 | Hydrolase | alpha/beta fold hydrolase [Actinocrispum wychmicini] | 67.14 | WP_165960812.1 |
| *orf29* | 338 | Regulator | LacI family transcriptional regulator [Saccharothrix ecbatanensis] | 75.53 | WP_184919824.1 |
| *orf30* | 617 | Glycosyl hydrolase | RICIN domain-containing protein [Dactylosporangium siamense] | 75.69 | WP_239135971.1 |
| *orf31* | 109 | Monooxygenase | antibiotic biosynthesis monooxygenase [Cohnella phaseoli] | 52.38 | WP_116065611.1 |
| *orf32* | 272 | Regulator | AraC family transcriptional regulator [Vitiosangium sp. GDMCC 1.1324] | 56.87 | WP_108075772.1 |
| *orf33* | 1048 | Transporter | multidrug efflux RND transporter permease subunit [Hyalangium sp. H56D21] | 79.13 | WP_224369323.1 |
| *orf34* | 376 | Transporter | efflux RND transporter periplasmic adaptor subunit [Myxococcus stipitatus] | 61.32 | WP_234065076.1 |
| *orf35* | 139 | Regulator | Transcriptional regulator, MarR family [Stigmatella aurantiaca DW4/3-1] | 52.24 | ADO71972.1 |
| *orf36* | 590 | Regulator | hypothetical protein A3F84_21795 [Candidatus Handelsmanbacteria bacterium] | 46.05 | OGG54796.1 |
| *orf37* | 172 | Unknown | hypothetical protein [Stigmatella hybrida] | 53.51 | WP_225414223.1 |
| *orf38* | 365 | Unknown | hypothetical protein E6G27_00285 [Actinobacteria bacterium] | 42.08 | TML43930.1 |

**TABLE S3. Proposed functions of the genes in the ZYM302 gene cluster and their closest relatives.**

#### TABLE S4

| pos. | $\delta_C$, type | $\delta_H$ (J in Hz) | COSY | HMBC | pos. | $\delta_C$, type | $\delta_H$ (J in Hz) | COSY | HMBC |
|---|---|---|---|---|---|---|---|---|---|
| Tyr-1 | | | | | Tyr-2 | | | | |
| 1 | 172.5, C | | | | 20 | 171.9, C | | | |
| 2 | 64.9, CH | 4.38, dd (10.6, 2.9) | 3 | 1, N-$CH_3$ | 21 | 66.7, CH | 3.70, dd (11.0, 2.8) | 22 | 20, 22, N-$CH_3$ |
| 3 | 27.0, $CH_2$ | a: 3.11, obs. | 2, 3b | 2, 5 | 22 | 27.5, $CH_2$ | a: 3.11, obs. | 21, 22b | 21, 23 |
| | | b: 2.79, obs. | 2, 3a | 2, 5 | | | b: 2.74, obs. | 21, 22a | 21, 23 |
| 4 | 131.0, $C^b$ | | | | 23 | 131.1, $C^b$ | | | |
| 5 | 130.8, $CH^b$ | 7.01-7.06, obs. | 6 | | 24 | 130.8, $CH^b$ | 7.01-7.06, obs. | 25 | |
| 6 | 115.0, $CH^b$ | 6.67, obs. | 5 | | 25 | 115.2, $CH^b$ | 6.67, obs. | 24 | |
| 7 | 154.1, $C^b$ | | | | 26 | 154.3, $C^b$ | | | |
| 8 | 115.0, $CH^b$ | 6.67, obs. | 9 | | 27 | 115.2, $CH^b$ | 6.67, obs. | 28 | |
| 9 | 130.8, $CH^b$ | 7.01-7.06, obs. | 8 | | 28 | 130.8, $CH^b$ | 7.01-7.06, obs. | 27 | |
| N-$CH_3$ | 41.0, $CH_3$ | 2.39, s | | 2, N-$CH_3$ | N-$CH_3$ | 41.5, $CH_3$ | 2.44, s | | 21, N-$CH_3$ |
| N-$CH_3$ | 41.0, $CH_3$ | 2.39, s | | 2, N-$CH_3$ | N-$CH_3$ | 41.5, $CH_3$ | 2.44, s | | 21, N-$CH_3$ |
| Thr-1 | | | | | Thr-2 | | | | |
| 10 | 168.1, C | | | | 29 | 167.7, C | | | |
| 11 | 53.4, CH | 5.16, d (4.9) | 12 | 1, 10, 12, 13, N-$CH_3$ | 30 | 57.0, CH | 5.14, d (1.9) | 31 | 20, 29, N-$CH_3$ |
| 12 | 74.1, CH | 4.00, m | 11, 13 | | 31 | 71.4, CH | 5.30, m | 30, 32 | 14, 32 |
| 13 | 14.9, $CH_3$ | 1.21, d (6.6) | 12 | 11, 12 | 32 | 16.8, $CH_3$ | 1.16, d (6.6) | 31 | 30, 31 |
| N-$CH_3$ | 31.0, $CH_3$ | 2.76, s | | 1, 11 | N-$CH_3$ | 34.0, $CH_3$ | 3.06, s | | 20, 30 |
| Leu-1 | | | | | Leu-2 | | | | |
| 14 | 168.8, C | | | | 33 | 170.6, C | | | |
| 15 | 52.2, CH | 5.33, dd (10.7, 5.6) | 15 | 14, 16, N-$CH_3$ | 34 | 61.7, CH | 3.12, obs. | 35 | 33 |
| 16 | 36.0, $CH_2$ | a: 1.79, m | 15, 16b, 17 | 14, 15, 17 | 35 | 38.7, $CH_2$ | a: 2.09, m | 34, 35b, 36 | 33, 36, 37, 38 |
| | | b: 1.39, obs. | 15, 16a, 17 | | | | b: 1.39, obs. | | |
| 17 | 24.5, CH | 1.32, obs. | 16, 18, 19 | | 36 | 25.4, CH | 1.62, obs. | 35, 37, 38 | 35, 37, 38 |
| 18 | 21.7, $CH_3$ | 0.82, d (6.6) | 17 | 16, 17, 19 | 37 | 22.8, $CH_3$ | 0.88, d (6.6) | 36 | 35, 36, 38 |
| 19 | 23.5, $CH_3$ | 0.91, d (6.6) | 17 | 16, 17, 18 | 38 | 22.7, $CH_3$ | 0.91, d (6.6) | 36 | 35, 36, 37 |
| N-$CH_3$ | 29.1, $CH_3$ | 2.98, s | | 10, 14 | N-$CH_3$ | 35.9, $CH_3$ | 2.01, s | | 29, 34 |

**Table S4. NMR data for metapeptin A**

TABLE S5

| Inhibitor | Inhibition, IC50 (μM) | | | |
|---|---|---|---|---|
| Inhibitor | hsMetAP1 | hsMetAP2 | EcMetAP | mtpF |
| Bengamide B | 29* | 18* | 0% inh. @ 2 mM | 0% inh. @ 2 mM |
| TNP-470 | 0% inh. @ 2 mM | 0.69 ± 0.05 | 0% inh. @ 2 mM | 0% inh. @ 2 mM |
| Metapeptin A | 0% inh. @ 2 mM | 0% inh. @ 2 mM | 0% inh. @ 2 mM | 0% inh. @ 2 mM |
| Metapeptin B | 49.71 ± 3.98 | 0% inh. @ 2 mM | 0.47 ± 0.29 | 0% inh. @ 2 mM |
| NMe-Monomer | 0% inh. @ 2 mM | 0% inh. @ 2 mM | - | - |

**Table S5.** S**ummary of MetAP enzymatic assay inhibition kinetics against known inhibitors, metapeptin A, metapeptin B, and NMe-Monomer. ***IC50 data for bengamide B taken from Garcia-Ruiz and Sarabia (49).

**Supplementary Text**

Metapeptin B structure elucidation

The UV/vis and NMR spectroscopic data for the compound in $CD_3OD$ displayed typical features of tyrosine-containing peptides. The two-dimensional NMR analysis revealed the amino acids components (Figure S4, Table S4).

We first investigated the two doublets in the aromatic region at δH 6.67 and 7.03 which comprise an AAXX system, and a key HMBC correlation between the aliphatic protons (δH 2.79 and 3.11, δC 27.0) indicated a tyrosine component.

Secondly, a methyl group (δH 1.21, d, *J*=6.6, δC 14.9) was then selected as a start point for COSY analysis. The COSY correlation between the $CH_3$ and a methine at δH 4.00, and HMBC revealed a further methine δH 5.16. The independent spin system of $XCHCHCH_3$ indicated a threonine moiety.

Next, COSY experiment revealed that two methyl groups (δH 0.82, d, J=6.6, and 0.91, d, *J*=6.6) were linked via a CH at δH 1.32 which is adjacent to a $CH_2$ (δH 1.39 and 1.79). Further COSY and HMBC showed an independent spin system of $XCHCH_2CH(CH_3)_2$ indicating the presence of a Leucine.

The signals resonating at δH 2.39 (6H, s), 2.76 (3H, s) and 2.98 (3H, s) with key HMBC correlations to the a protons as well as carbonyl groups indicated *N*-methyl or *N,N*-dimethyl, and revealed the order of the tri-peptide monomer as Tyr-Thr-Leu.

Similarly, a second monomer fragment was identified. The key HMBC correlations of β-H on threonine (δH 5.30) to the carbonyl group of Leu (δC 168.8) suggested that the two monomers were conjugated via ester bonds.
